# Assessing the potential antifungal resistance risk from dual use of a mode of action in agriculture and medical treatment of human pathogens

**DOI:** 10.1101/2024.05.21.595086

**Authors:** Neil Paveley, Frank van den Bosch, Michael Grimmer

## Abstract

A mechanistic basis is described for assessment of resistance risk to medical anti-fungal treatments from agricultural use of fungicides of the same mode of action. The following need to occur in landscape environments for a risk to be posed by ‘dual use’: (i) emergence, whereby a resistant strain emerges by mutation and invasion, (ii) selection, whereby a mutation conferring a fitness advantage is selected for in the presence of fungicide, and (iii) exposure of humans to resistant strains from the landscape, potentially resulting in invasive fungal infection (IFI). We identify 20 human fungal pathogens for which there is evidence that all three processes above could, in principle, occur. A model is derived for quantitative analysis to explore what determines resistance emergence and selection in human pathogens in landscape environments. Emergence and selection were particularly affected by fitness cost associated with the resistance mutation(s) and fungicide concentration.

Emergence was also determined by the amount of pathogen reproduction (related to pathogen population size). The findings were related to an example case of observational data from the Netherlands for Aspergillus fumigatus. The analysis supports previous work that compost, including bulb waste, is towards the high-risk end of the spectrum for this species. Agricultural soils, non-agricultural land and grassland were lower risk. More generally, across species, the model output suggests that if fungicide resistance is associated with even a small fitness cost, then environments with low fungicide concentrations, such as field soils and semi-natural environments (e.g. woodland), may not be conducive to resistance emergence or selection.

## INTRODUCTION

The case of azole resistant *Aspergillus fumigatus* (ARAf) has highlighted that agricultural use of fungicides can potentially contribute to resistance development in non-target species that are also human pathogens; with consequences for human health. A balance is needed between maintaining availability of fungicides which are important for food security (Steinberg & Gurr, 2020) and protecting the efficacy of antifungals used to treat serious invasive fungal infections (IFI) (Robbins et al., 2017). Risk arises where active substances with the same mode of action (MoA) are used to control fungal pathogens of crops and to treat IFI in medicine (Bastos et al., 2021; Fisher et al., 2022). The current parallel development of fungicides and antifungals targeting dihydroorotate dehydrogenase (DHODH) for use in both agriculture and medicine (Verweij et al., 2022) has highlighted the need for a method to determine whether a new agricultural use poses a material risk to medical use and, if so, whether practical mitigation measures can reduce that risk to an acceptable level. Ideally, such a risk assessment method should be broadly applicable across MoAs and across fungi which are human pathogens and are potentially exposed to fungicides in the agricultural environment. To be of maximum value, it should also require only information that can be obtained during the development of new active substances – rather than being a *post-hoc* assessment after product approval.

The World Health Organisation (WHO) has listed human fungal priority pathogens and categorised them as ‘critical’, ‘high’ or ‘medium’ priority (WHO, 2022). Human fungal pathogens were also reviewed by Köhler et al. (2015) and Gisi (2022). There are differences in the fungi included in the lists from these three sources, partly resulting from the different focus of each of the papers. In some cases fungi are listed at genus level, despite substantial differences between species in aspects of their lifecycle that may be relevant to risk. We reconcile these three sources and identify pathogens of concern at a species level.

The resulting list of priority human pathogenic fungi may contain species for which agricultural use of fungicides could, in principle, pose a risk to medicine and other species for which a material risk is mechanistically improbable. A key role of risk assessment is to discriminate between these two groups of fungi. Section One of the work was, therefore, to define the epidemiological and evolutionary mechanisms that need to occur to create a potential risk, and then review the literature to determine for each pathogen whether each of the relevant mechanisms can occur.

For pathogens where all the mechanisms could, in principle, co-occur to create a potential risk, attention then turns to the degree of risk. There is likely to be a distribution of risk from different species and environmental compartments. The concept of ‘hotspots’ (that pose a potential risk to human health) and ‘coldspots’ (that do not) has been introduced, as an approach to identifying combinations of fungus, active substance and environment where agricultural use poses a risk (Gisi, 2022). If agricultural use of a new fungicide does not create a ‘hotspot’ then it could be argued that dual use of a MoA in both agriculture and medicine is acceptably safe. Note that although the risk results from dual use of the same MoA, the assessment of risk needs to include consideration of the intrinsic activity of the active substance (a.s.), as different a.s. within a MoA can have substantially different activity against particular human pathogen species. Although the concept of ‘hot’ and ‘cold’ spots appears binary, the figures in Gisi (2022) show the continuum of risk factors that were considered, resulting in an assessment of whether the particular combination under consideration is towards the high- or low-risk end of the spectrum. Section Two of the work reported here was to develop a method for quantitative analysis, to explore this risk spectrum further. We addressed the question: What are the key determinants of resistance risk and, hence, what determines whether an active substance/environmental compartment combination constitutes a hotspot for a particular pathogen? Furthermore, can we identify aspects of the system that may be sensitive to mitigation measures to reduce risk? This article focusses on agricultural use of fungicides and the resulting potential for exposure of humans to resistant strains arising from the landscape. Assessment of the resistance risk from antifungal use in medical settings is not addressed. The analysis considered risks from specific environmental compartments in the landscape. Although the exposure of a human pathogen to an agricultural fungicide originates predominantly from treatments targeting crop pathogens in agricultural fields, there may be flow of substrate, pathogen and fungicide between environments. Mechanisms for such flow include, for example: transfer of plant material for composting; movement of mycelium, resting bodies or spores; and movement of fungicide as residues in plant material or by spray drift to adjacent land. Adjacent land may include ‘semi-natural’ environments, which we define here as areas (such as woodland) which are not agricultural or built environments.

The rationale for the analysis of fungicide resistance evolution in human fungal pathogens in landscape environments is based on understanding gained from experimental and modelling studies on the effects of fungicides on resistance evolution in target fungal crop pathogens in agricultural fields.

Two evolutionary phases have been defined (van den Bosch et al., 2011) which need to occur for a risk of resistance to be present:

Emergence phase: A resistant strain emerges by mutation and invasion.

Selection phase: A mutation which confers a fitness advantage will be selected for in the presence of fungicide, resulting in an increase in the proportion of the population carrying the mutation.

The combination of the emergence and selection phases results in the potential for exposure of humans to resistant strains, resulting in IFI. The degree of exposure depends on fungal population density, frequency of resistance and whether humans (at times they are susceptible to IFI) are exposed to fungal propagules coming from the landscape.

For agricultural fungicide use to create a risk to medicine, emergence, selection and human exposure all need to occur. These three processes underpinned Section One of the work reported here; defining the mechanisms that need to occur to create a potential risk, and determining for each human pathogen of concern whether all the relevant mechanisms could, in principle, occur.

For pathogens that represent a potential risk, the practical need is for risk assessment to support decisions on product development and mitigation measures. The aim of those decisions is to ensure that the probability of human infection by a resistant fungal strain (arising from the landscape) remains below an acceptable threshold for as long as possible. This requires that the emergence time is long and the selection rate small.

The quantitative analysis reported in Section Two used a modelling approach to explore the question: what determines the emergence time and selection rate of resistance to agricultural fungicides in a human pathogen exposed to fungicides in a landscape environment? The approach was based on models tested and used previously to guide resistance management options for crop pathogens (van den Bosch et al., 2015).

Experience with crop pathogens has shown that a combination of experimentation and modelling can support resistance management decisions. Modelling cannot yet predict timescales for particular outcomes accurately but it can identify key determinants of those outcomes and the direction of their effects. Factors and variables explored, for the specific landscape environments under consideration, were aspects of the physical environment, biological environment, fungal characteristics and fungicide efficacy.

Lastly, we apply the model predictions for risk determinants to an example case of *A. fumigatus* in agricultural fields, compost (including bulb waste) and semi-natural environments in The Netherlands. We review the extent to which the model outputs reinforce, or add to, existing understanding.

### MECHANISMS NECESSARY FOR RESISTANCE EVOLUTION

This section outlines mechanisms of fungicide resistance evolution which are relevant for risk assessment. Qualitative risk assessment should determine whether all the necessary mechanisms for resistance can or cannot occur for a given pathogen and landscape environment. Quantitative risk assessment should be based on a quantitative representation of the mechanisms.

### Emergence phase

When resistance is not present in the genetic background of the population, it has to arise through mutation and invasion. Mutation refers to one or more heritable changes in individual cells conferring some level of resistance. Invasion refers to the increase in frequency – by selection or genetic drift – to a sufficiently high number that the resistant strain is unlikely to die out by random chance. We are not, in this analysis, concerned with which specific genetic mechanisms cause the resistance to arise (target site mutation, upregulation, enhanced efflux, etc), but rather with the resulting sensitivity phenotype that drives evolution.

At the start of the emergence phase a fungal population consists of sensitive strains. Occasionally a mutant spore carrying the resistance mutation will arise. There is a high probability that this spore, or the descendants of this spore, will die out because the vast majority of spores die, rather than form infectious lesions. Hence, it is likely to take many such mutation events before the resistant strain emerges in the population (Figure 1).

**Figure 1.**
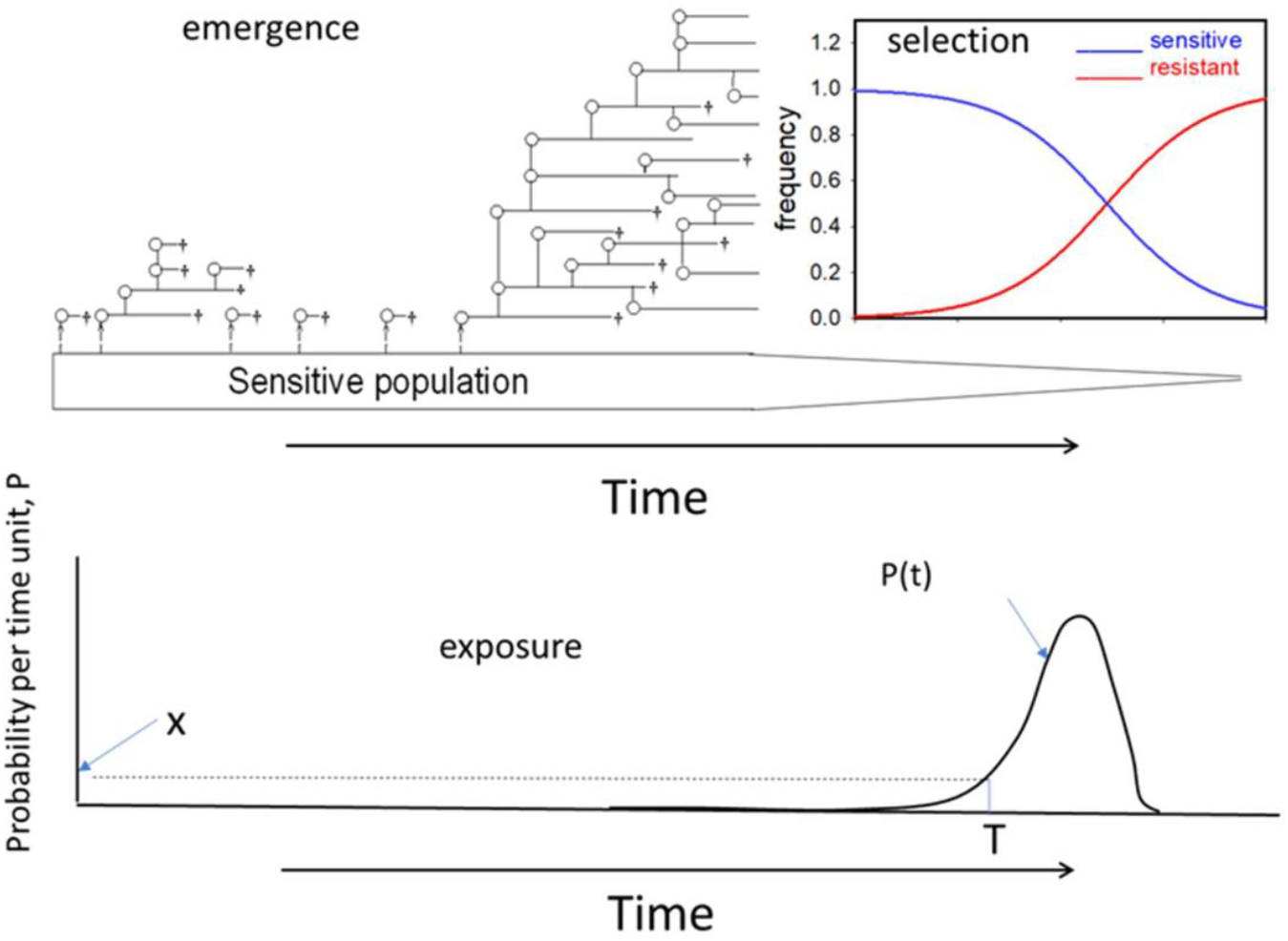
Illustration of the emergence and selection phases and resulting probability for exposure of humans to a fungicide resistant strain.

Modelling of plant pathogens shows that emergence may happen, by random chance, prior to introduction of a new fungicide, but is much more likely to occur after introduction (Hobbelen et al., 2014) because of the fitness of the resistant mutant in the presence of fungicide. The time period from introduction to emergence depends, among other things, on the number of mutant spores developing per time unit, which in itself depends on the size of the sensitive population. The time to emergence also depends on the transmission rates of the resistant lesion. Emergence is thus governed, to a major extent, by the basic reproduction number, R0 (the number of daughter lesions per mother lesion in a fixed environment; Diekmann et al., 1990), of the resistant strain in the sensitive population. Note that, in many applications, R0 is defined as the number of offspring where the pathogen is not present (or is present at infinitesimally low density). In the resistance case we are interested in the number of offspring of a resistant lesion in a population of sensitive lesions.

### Selection phase

In the selection phase, the *per capita* rate of increase, *r*, of the fungicide-resistant strain is larger than that of the sensitive strain (Figure 1), due to the greater effect of the fungicide on the sensitive strain. This increases the fraction of the pathogen population that is resistant to the fungicide. It is important to note that the effect of selection is to change the proportion of the resistant strain in the total population. Selection is not about absolute numbers. In the selection phase the governing quantity is the rate of increase, *r*, of the resistant strain (relative to that of the sensitive population) and not, as in the emergence phase, the absolute number of offspring.

### Exposure of humans

Although resistant propagules may be released during the emergence phase, the absolute number of resistant spores is very small and localised, so the risk of exposure of humans to the resistant strain is likely to be negligible. During the selection phase the probability (P) for exposure increases, initially exponentially (Figure 1) (Crow & Kimura, 1970). At a certain time (T) the level of exposure will exceed a threshold value (X), beyond which the level of risk is considered to be unacceptable. The aim is to delay T, ideally indefinitely.

### Processes determining resistance emergence, selection and exposure in landscape environments

Emergence, selection and exposure may occur in a range of environments in the landscape. For this analysis we consider three environmental compartments: agricultural fields, compost from plant waste, and non-agricultural environments where fungicides are seldom used; mainly grassland for grazing livestock and semi-natural environments (such as woodland). For brevity we include grassland within the general term ‘semi-natural environments’. Humans may be exposed to propagules of resistant strains emitted from these environments, thus affecting the efficacy of antifungals prescribed in medical settings to treat IFI (see schematic Figure 2). The analysis here is limited to evolutionary processes occurring in the three landscape environments, i.e. to the left of the dotted line in Figure 2.

**Figure 2.**
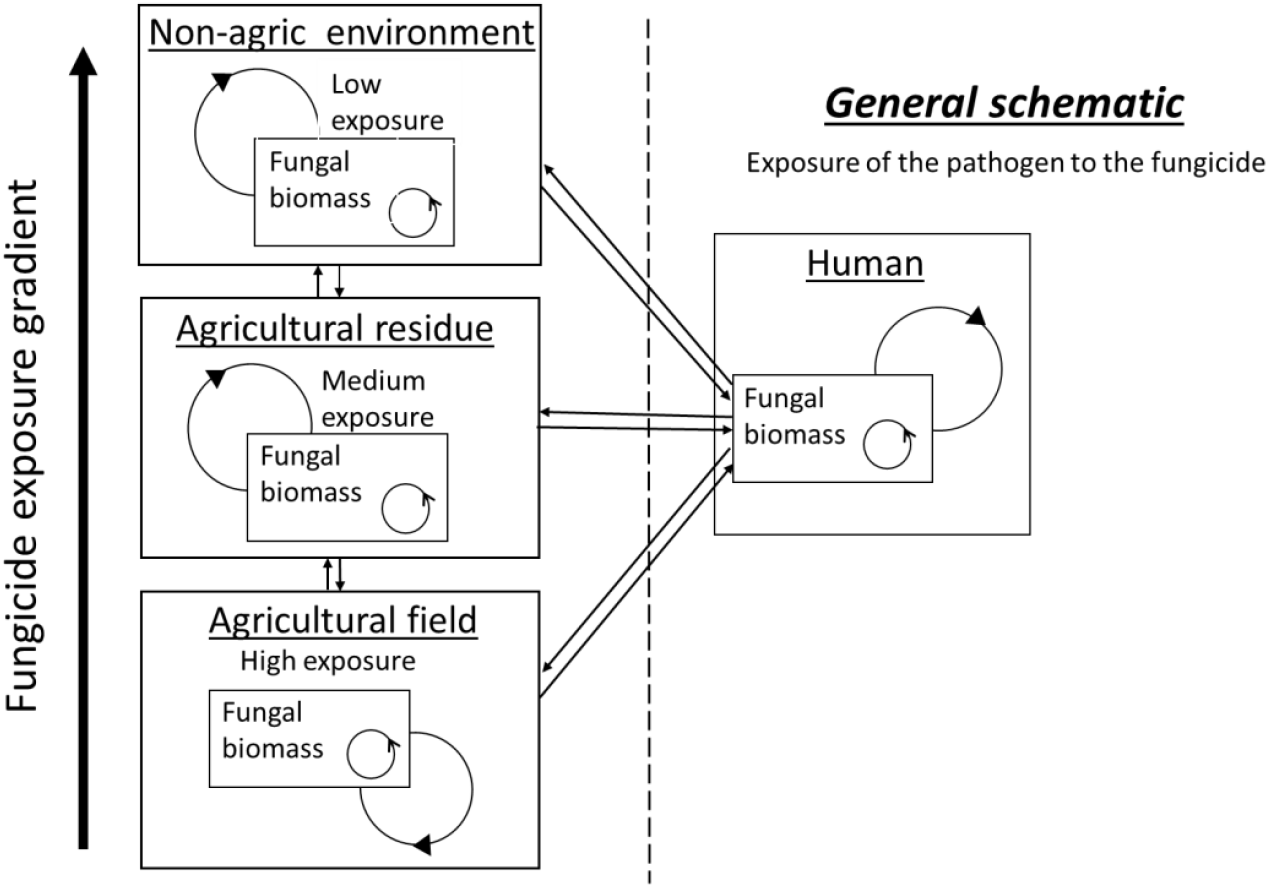
Schematic of the three environmental compartments under consideration within the landscape (left of dotted line) and the linkages between them (arrows) representing flow of one or more of substrate, pathogen and fungicide residues. Medical settings, to the right of the dotted line, are out of scope of this article. As presented this schematic is for a saprotrophic pathogen, but is easily adjusted to include other environmental compartments. The agricultural waste compartment is that waste that can support populations of the pathogen under consideration. For example for *A.fumigatus* this includes compost heaps but not anaerobic degradation.

The arrows in Figure 2 represent coupling between environments caused by flow of material. To be generic, the schematic shows the possible bidirectional flow between all the environments. Particular directions of flow may or may not apply to particular pathogens or landscape systems. Flows which are likely to be relevant in many cases include, for example: movement of plant material (substrate for fungal growth) from agricultural fields to composted plant waste (which may also transfer the pathogen and fungicide residues); transfer of fungal propagules (for example by airborne spores) between landscape environments and from the landscape to humans; and movement of fungicides in the environment (for example by spray drift to land adjacent to fields).

The effect of a fungicide on fungal biomass growth will differ between resistant and sensitive strains, thus creating evolutionary pressure for emergence and selection. The scale of the evolutionary pressure will depend on the concentration of the fungicide present in each landscape environment and the intrinsic activity of the active substance. The concentration will be highest in the crop plants to which the fungicide is directly applied, lower in composted plant waste, and lower again in non-agricultural environments. The effects of fungicide concentrations that are substantially below the dose applied to crops need to be understood, because of concern about the effects of ‘sub-lethal doses’ driving resistance.

Within each of the three landscape environments and across different pathogen species, certain processes of fungal growth will occur; illustrated as a generic schematic in Figure 3, based on the work of Lamour et al. (2000, 2002), who considered the growth a fungal population in relation to the dynamics of carbon and nitrogen. The extent to which particular substrates support the growth of particular fungi will differ according to the characteristics of the substrate and the methods fungi have evolved to extract carbon, nitrogen and other nutrients. The mechanisms of reproduction and colonisation will also differ between fungi.

**Figure 3.**
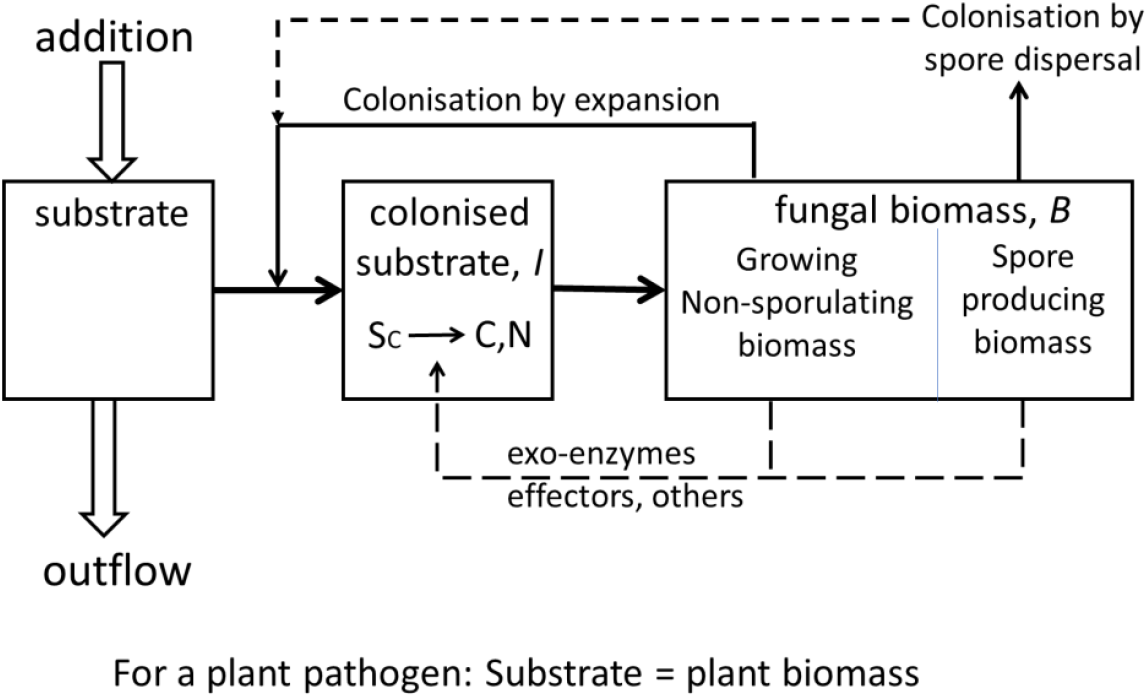
Schematic for growth of a fungal population colonising a substrate in a specific environmental compartment by biomass growth and by colonisation of new biomass via sporulation. Variables are: substrate density (*S*); *c*olonised substrate density (I); carbon source density, carbon freed up from the colonised substrate in a form that the fungus can take up (*C*); fungal biomass density (*B*). N is the nitrogen concentration, which we do not further consider in the model.

Section One below (qualitative risk analysis) uses the structures shown in Figures 2 and 3 to guide a review of the literature. The aim is to identify the human pathogen species for which there is evidence that all the mechanistic processes can occur that are necessary for resistance to emerge and be selected for in the landscape and pose a potential risk to medicine through exposure of humans to resistant strains.

Section Two below (quantitative risk analysis) uses the structures in Figures 2 and 3 to derive a quantitative model to explore determinants of emergence and selection.

## SECTION ONE: IDENTIFYING HUMAN PATHOGENS WITH A POTENTIAL MECHANISTIC ROUTE FOR RESISTANCE TO EVOLVE IN THE LANDSCAPE AND TRANSFER TO MEDICAL SETTINGS

### Methods

The primary information source for identifying the most important human fungal pathogens was the World Health Organization (WHO) fungal priority pathogens list to guide research, development and public health action (WHO, 2022). The list focused on fungal pathogens that can cause invasive systemic fungal infections for which drug resistance or other treatment and management challenges exist. The pathogens included were ranked, then categorized by the WHO into three priority groups (critical, high, and medium). In most cases, the pathogens were referred to down to species level, however for some pathogens only the genus was provided in the WHO report. In such cases, two further sources were used to determine the species of importance, based on the genera reported by WHO. These were from the first table in Gisi (2022), which was reportedly based on information from Pfaller & Diekema (2004), and Köhler et al. (2015).

For each of the human fungal pathogens identified from those sources, information on their landscape occurrence, reproduction and transmission from the landscape to humans was sought from the literature and the evidence summarized. The results of this literature search for information are given in Supporting information S1, together with the sources.

The landscape was mainly taken to include agricultural fields, plant waste and semi-natural environments. Definitions of substrates within those environments are given in the literature cited.

The summarized evidence on landscape occurrence was used to assign each of the human pathogens to a category (high, medium, low or not reported) or a mixed category (e.g. medium / high) where a single category did not represent the range or uncertainty of information in the literature. Whilst the evidence on occurrence was broad, and included anthropogenic settings such as households and hospitals, the categorization was performed focusing on occurrence in landscape environments and substrates.

Occurrence in an environment or substrate does not necessarily indicate pathogen growth and reproduction in that environment – pathogen material may have been deposited there, for example as spores produced elsewhere. Growth and reproduction are necessary for resistance evolution, and hence for potential resistance risk. We assumed that pathogens that accumulate to a significant density in natural environments also grow in those environments.

The summarized evidence on environmental reproduction was used to assign each of the human pathogens to a category depending on propagule or spore type referred to in the literature (asexual, sexual and asexual or not reported). In some cases the evidence for presence of a particular propagule or spore type was from *in vitro* observations rather than from observations in the natural environment. This was considered a plausible approach that reflected the potential for propagules and spore types to exist in the environment even if they had not been directly observed.

Evidence on human infection was used to categorize the landscape transmission to humans as high, low or not reported. In the case of pathogens in the landscape, the fungus may be transported into the human body via airborne or water-borne propagules or spores, for example. In the case of pathogens that rely on human-to-human transmission for infection, this transmission could occur variously by airborne spores, medical devices, surfaces in a home or by direct human-to-human contact. Rather than having regard to the various mechanisms by which the pathogen may infect humans, the categorization of landscape transmission was carried out based on the extent to which the landscape is considered to represent a source of inoculum for human infection.

## Results

Analysis of the published sources from Gisi (2022), Köhler et al. (2015), Pfaller & Diekema (2004) and WHO (2022), resulted in 27 important human fungal pathogens being identified to species level (Table 1), belonging to 16 different genera.

**Table 1.**
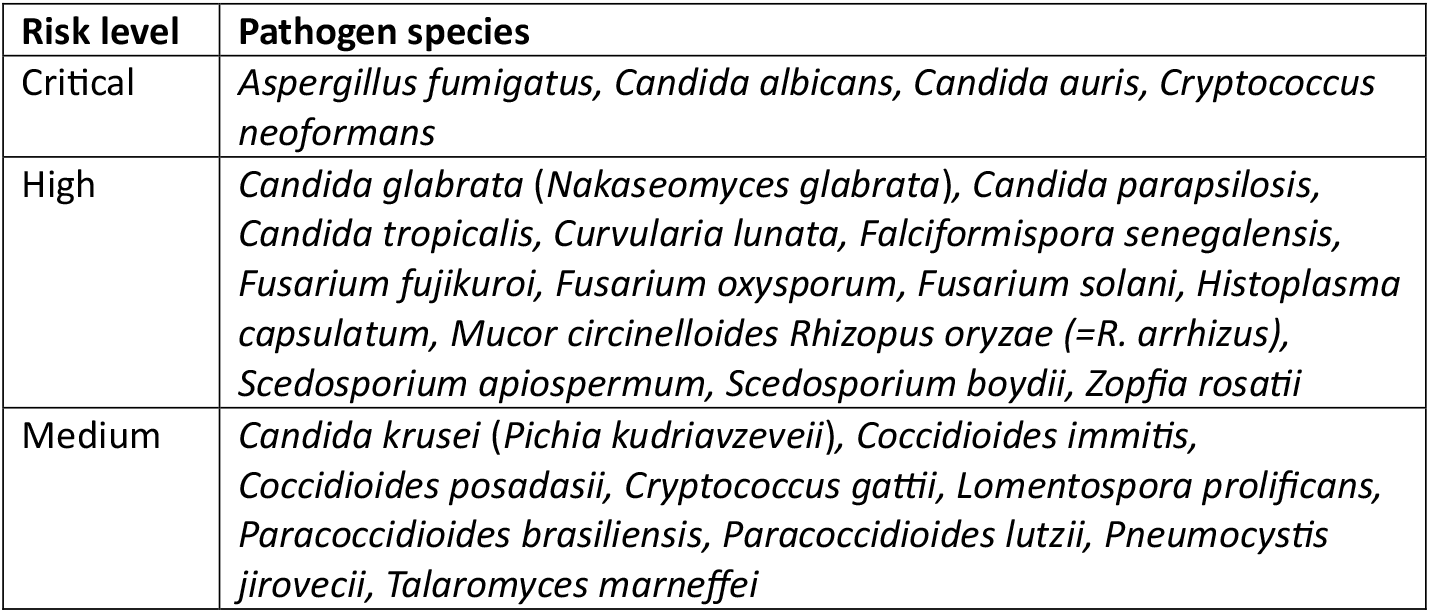
Human fungal pathogens species of concern. Risk categories are from the World Health Organization (WHO, 2022)

### Occurrence in landscape environments

Two of the four WHO ‘critical priority’ pathogens – *Aspergillus fumigatus* and *Cryptococcus neoformans* – were determined to have high reported landscape occurrence. Four of the fourteen ‘high priority’ pathogens and one of the nine ‘medium priority’ pathogens were determined to have high / medium landscape occurrence. Two of the four WHO ‘critical priority’ pathogens – *Candida albicans* and *Candida auris* – were determined to have low landscape occurrence. Two of the fourteen ‘high priority’ pathogens and one of the nine ‘medium priority’ pathogens were determined to have low or no reported landscape occurrence (Table 2).

**Table 2.**
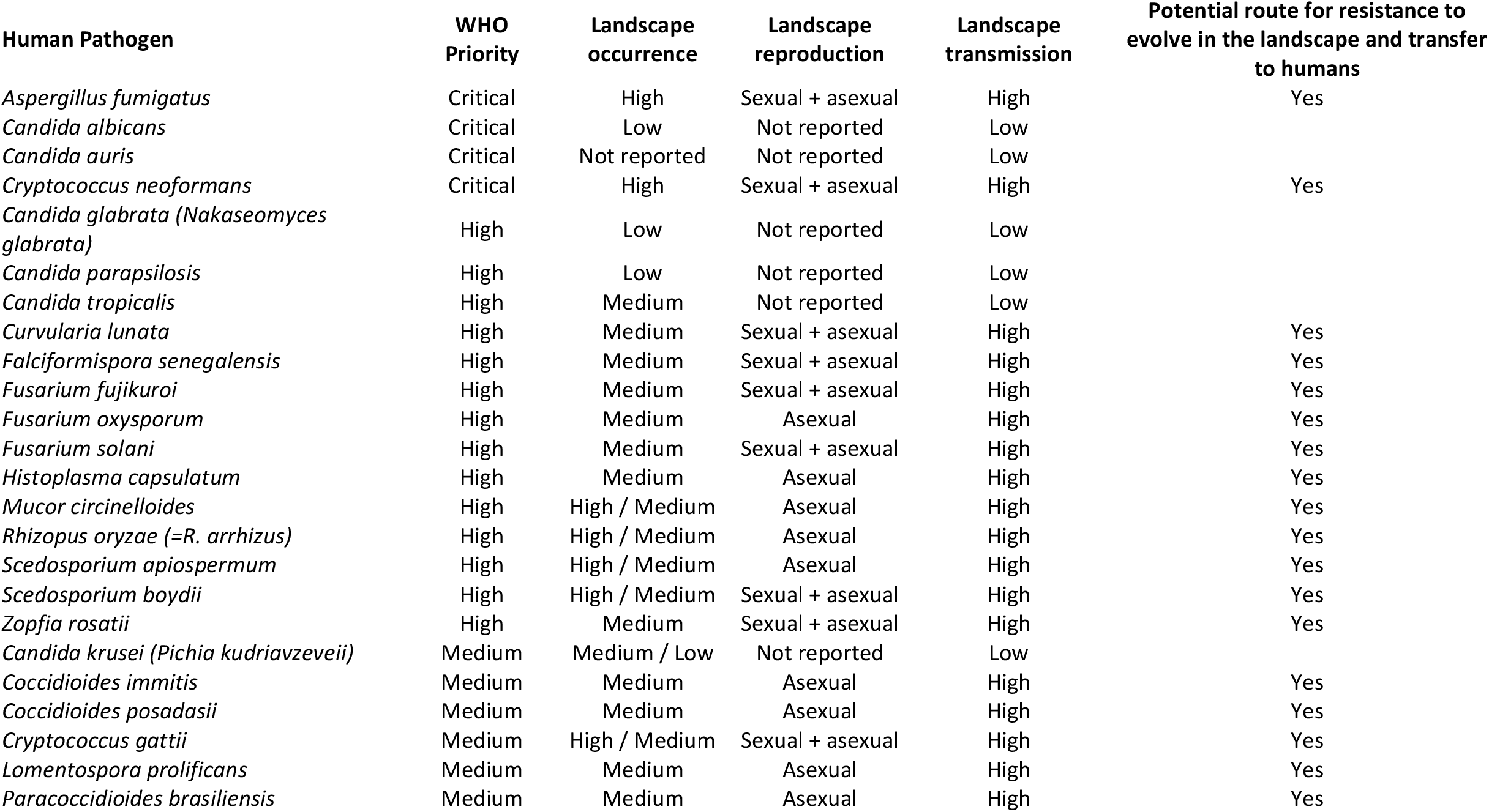
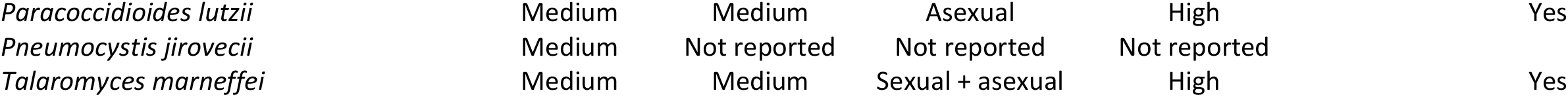
Human fungal pathogen species and their and potential mechanistic route for resistance from agronomic environments to medical settings.

### Reproduction

Of the four WHO ‘critical priority’ pathogens, two were categorized as sexual and asexual and two were categorized as having no reported landscape reproduction. Of the fourteen ‘high risk’ pathogens, six were categorized as having sexual and asexual spores, five as having asexual spores only and three as having no reported landscape reproduction. Of the nine ‘medium risk’ pathogens, two were categorized as having sexual and asexual spores, five as having asexual spores only and two as having no reported landscape reproduction (Table 2). In all cases, species which do not have reported reproduction in the landscape are those for which the primary infection route to humans is believed to be by human-to-human spread.

### Landscape transmission to humans

Of the 27 priority fungal pathogens, 20 were determined to have the landscape as an important potential source of human infection. Landscape transmission was determined to be of lower importance for 6 of the pathogens; these were all *Candida* species (including critical, high and medium priority pathogens). The remaining, *Pneumocystis jirovecii* (medium priority), has not been reported to have transmission from the landscape to humans.

The information given in Supporting information S1 enables the general schematic flow diagram (Figure 2) to be re-drawn showing the processes and couplings relevant to an individual pathogen species; as shown in Figure 4 for the example of *A. fumigatus*. The unidirectional arrows in the centre of Figure 4 represent the flow of fungal inoculum from different environmental compartments to humans.

**Figure 4.** Schematic for growth of an *A. fumigatus* population colonising a substrate by biomass growth and by colonisation of new biomass via sporulation.

The 7 pathogens not considered to have a mechanistic route for spread of resistance from agronomic to medical settings were the six *Candida* species and *Pneumocystis jirovecii*.

Proving the negative is difficult, so the presence of some potential risk to medicine from agricultural fungicide use cannot be discounted completely by this analysis. Some of these 7 pathogens are believed to occur in the natural environment to some limited extent.

However, any potential risk of spread is likely to be low.

Pathogens with a medium or high landscape occurrence, along with landscape reproduction and transmission from the landscape to humans, were considered to have the necessary mechanisms to allow, in principle, for resistance to evolve in landscape environments and transfer to humans. Of the 27 human fungal species listed in Table 1, 20 pathogens from 14 different genera were identified as having such a mechanistic route. These included 2 critical priority pathogens, 11 high priority pathogens and 7 medium priority pathogens.

The type of reproduction (sexual and asexual or asexual only) was not included here in the assessment of the presence of a mechanistic route. They indicate contrasting evolutionary potential that could be important in future work.

For the 20 pathogens found to have a potential mechanistic route for spread of resistance from agricultural environments to humans, the qualitative analysis has highlighted a potential risk. To assess the degree of risk, quantitative analysis is needed to determine the extent to which the pathogen is present in specific landscape environments and the degree of exposure to agricultural fungicides in those environments. This is explored in Section Two.

## SECTION TWO: TOWARDS QUANTITATIVE RISK ASSESSMENT

A model was developed for the human pathogen-agricultural fungicide system, to address the question: what determines the emergence time and selection rate of resistance to agricultural fungicides in a human pathogen? The model was used to explore the effect of parameters/variables representing aspects of the physical environment, biological environment, fungal characteristics and fungicide efficacy. Values were obtained from the literature for fungi in agricultural and other landscape environments, to enable initial estimates of parameter values. However, the published studies were not designed to enable such parameter estimation, so this analysis should be considered as a first iteration of modeling, which will help to identify key parameters and prioritise future data gathering.

To properly assess risk, coupling of environmental compartments is necessary to account for the flow of substrate, fungicide and pathogen between environments - for example, the flow of substrate, pathogen and fungicide from agricultural fields to composted agricultural waste. Further data are required to quantify such flows, so the analysis here considers an environmental compartment in isolation. The results should therefore be interpreted with caution.

In the sub-sections below (and Supporting Information) we describe: (i) model development, (ii) parameter estimation, (iii) model results, and (iv) interpretation of risk predictions from the models for an example case of *A. fumigatus* in the Netherlands.

### Model development

#### Representing fungal growth and colonisation

The model of Lamour et al. (2000, 2002) considered the growth of a saprotrophic fungal population in relation to the dynamics of carbon and nitrogen. For the purposes of this analysis, the fungus is exploiting a substrate which differs between environments, for example, organic material in soil, compost or crop plants. Given the sparse data available for parameter estimation, we modify the model by considering carbon only and with simplified dynamics, based on the schematic representing one environmental compartment (Figure 3).

The equations for the rate of change of these densities are For substrate density, S:

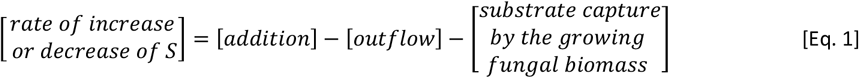

For the example of compost, addition of material in Eq. 1 represents the rate at which new plant material is added to the compost heap and outflow represents the removal of compost for further use.

For colonised substrate density, I:

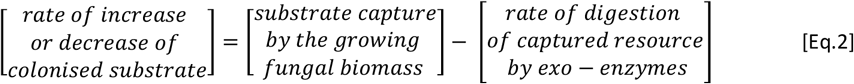

For the compost example, substrate capture in Eq. 2 represents the rate of colonisation of the plant material by a pathogenic fungus growing saprophytically, and rate of digestion is the rate of loss of colonised substrate due to enzymatic break down by the fungus.

For carbon freed up from the colonised substrate in a form available to the fungus:

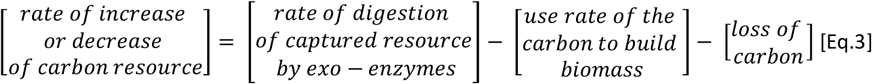

For compost, the rate of change of available carbon resource for the fungus in Eq. 3 will be affected by the rate of use of carbon to build fungal biomass of the pathogen and loss of carbon by other means (such as to other fungal and bacterial species).

For fungal biomass density, B:

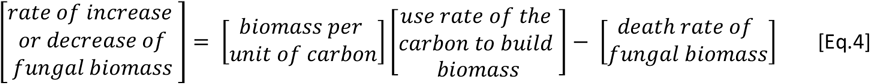

For the fungal pathogen growing saprophytically in compost, the rate of change of fungal biomass in Eq. 4 will be determined by the amount of biomass produced per unit of carbon acquired, the rate of use of available carbon and the death rate of the fungal biomass.

The processes described in equations 1 to 4 were formulated as a set of linked differential equations, describing rates of change. Equations were then derived for the *per capita* (intrinsic) rate of increase of the pathogen population, *r*, and the net-reproductive number, *R*_0_.

*r* is defined as the per capita increase in pathogen density per time unit. In population genetics, the fitness of a pathogen strain (Crow & Kimura, 1970) is defined as the intrinsic rate of increase, *r*, of that pathogen strain. *R*_*0*_ is defined in this context as the number of offspring per individual during its entire lifespan when availability of resource is not limiting.

The rate of selection for a resistant strain is determined by the difference in fitness, *r*, between the sensitive and resistant strain (Crow & Kimura 1970). Emergence of a resistant strain is related to *R*_*0*_. Applying the methods described by Mikaberidze et al. (2017) and the selection equation from Milgroom & Fry (1988), and incorporating the effects of fungicide dose, equations were derived which gave an analytical expression for emergence time, *P* (Eq.5) and selection rate, *s* (Eq.6). Full derivations of equations 5 and 6 are given in Supporting Information S2.

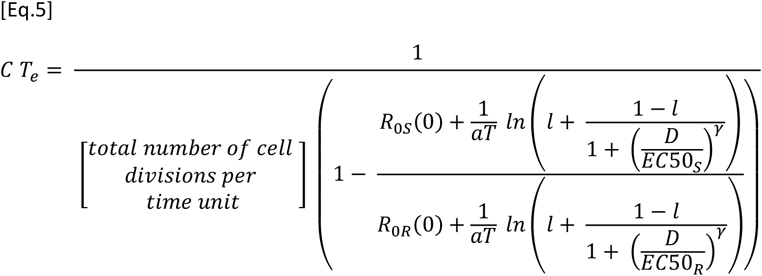

Parameters are R_0S_ (net reproduction number of sensitive strain), R_0R_ (net reproduction number of resistant strain), L (asymptote of dose-response), EC_50_S (EC_50_ sensitive strain) EC_50_R (resistant strain), aT (carbon absorption rate), and *γ* (dose response shape parameter). The emergence time *T*_*e*_ has a constant C in front of it (explained in Supporting Information S2) for which the numerical value is currently unknown, so we are limited to plotting lines of equal emergence time (see later figures) multiplied with an unknown constant. The ‘total number of cell divisions per time unit’ is also derived in terms of model parameters in Supporting Information S2.

Selection, *s*, is measured by *S* = *r*_*R*_ − *r*_*S*_ where *r*_*R*_ and *r*_*S*_ are the growth rates for the resistant and sensitive strains, respectively (Milgroom & Fry, 1988). For the model of a fungus in a given environmental compartment we get

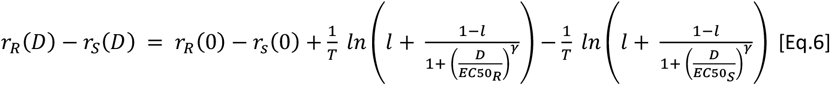

Parameter definitions as for Eq.5, above. When there is no fitness cost to being resistant *r*_R_=*r*_S_ in the absence of fungicide. If there is no fitness cost and the dose response curve has asymptote at zero (i.e. an infinite dose of fungicide would, in theory, reduce pathogen density to zero) and the resistance is absolute (i.e. the resistant strain is not controlled by the applied dose of fungicide), then 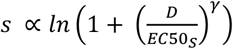

Data were available from the literature to estimate parameters for equations 5 and 6 only for a range of agriculturally relevant fungi, to enable a preliminary analysis of sensitivity of emergence time and selection rate to variation in each parameter.

### Parameter estimation

Parameter ranges and default values used for sensitivity analysis are given in Table 3 with sources. Although human fungal pathogens (particularly *A. fumigatus)* have been studied recently in landscape environments, data to estimate the required parameters remain sparse. Therefore, where no parameter values are known from published sources, we used the values known for plant pathogens. This especially holds for the net reproductive number of the pathogens.

**Table 3.**
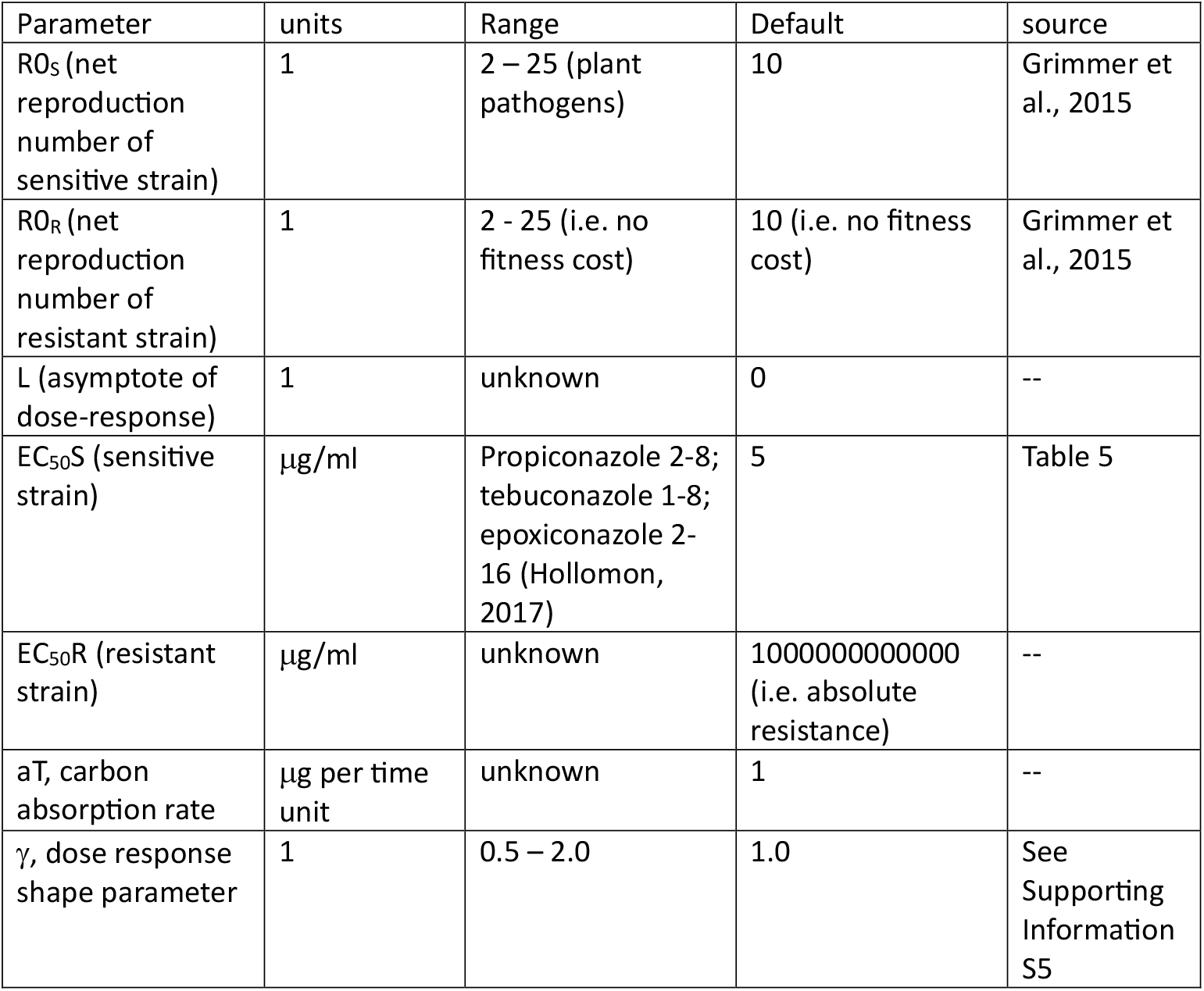
Parameter ranges and default values.

| Parameter | units | Range | Default | source |
| --- | --- | --- | --- | --- |
| R<sub>0S</sub> (net reproduction number of sensitive strain) | 1 | 2 – 25 (plant pathogens) | 10 | Grimmer et al., 2015 |
| R<sub>0R</sub> (net reproduction number of resistant strain) | 1 | 2 - 25 (i.e. no fitness cost) | 10 (i.e. no fitness cost) | Grimmer et al., 2015 |
| L (asymptote of dose-response) | 1 | unknown | 0 | -- |
| EC<sub>50S</sub> (sensitive strain) | µg/ml | Propiconazole 2-8; tebuconazole 1-8; epoxiconazole 2-16 (Hollomon, 2017) | 5 | Table 5 |
| EC<sub>50R</sub> (resistant strain) | µg/ml | unknown | 1000000000000 (i.e. absolute resistance) | -- |
| aT, carbon absorption rate | µg per time unit | unknown | 1 | -- |
| γ, dose response shape parameter | 1 | 0.5 – 2.0 | 1.0 | See Supporting Information S5 |

### Model results – determinants of emergence time and selection rate

#### Emergence time for fungicide resistant strain

Contour lines for values of the emergence time are shown in Figure 5. These are calculated from equation 5 and using the parameter values in Table 3. Note again that the three contour lines in Figure 5 show relative values, CT_emerge_; from top right to bottom left representing 1-fold, 10-fold and 100-fold increases in emergence time, respectively.

**Figure 5.**
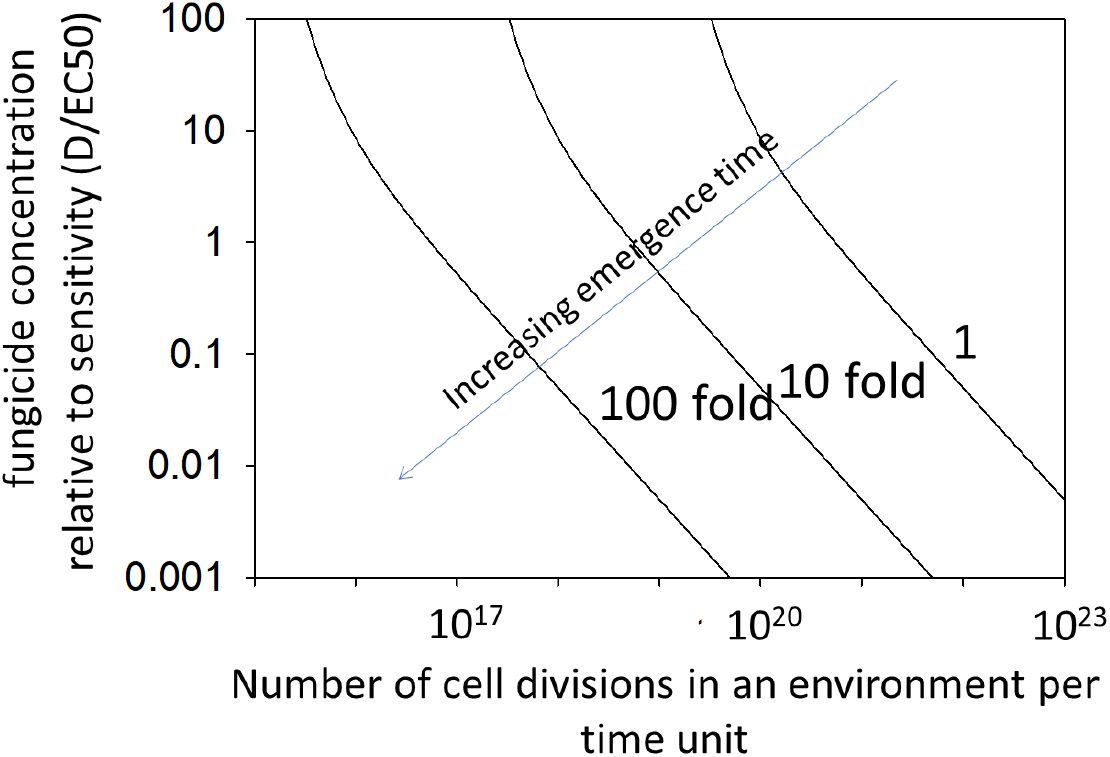
The contours of relative emergence time, as determined by pathogen reproduction and fungicide concentration. The three contour lines show order of magnitude increases in emergence time from top right to bottom left.

Changing one parameter at a time, across the range of parameter values given in Table 3 (excluding fitness cost), caused small or moderate changes in the emergence time contours (Figure 6).

**Figure 6.**
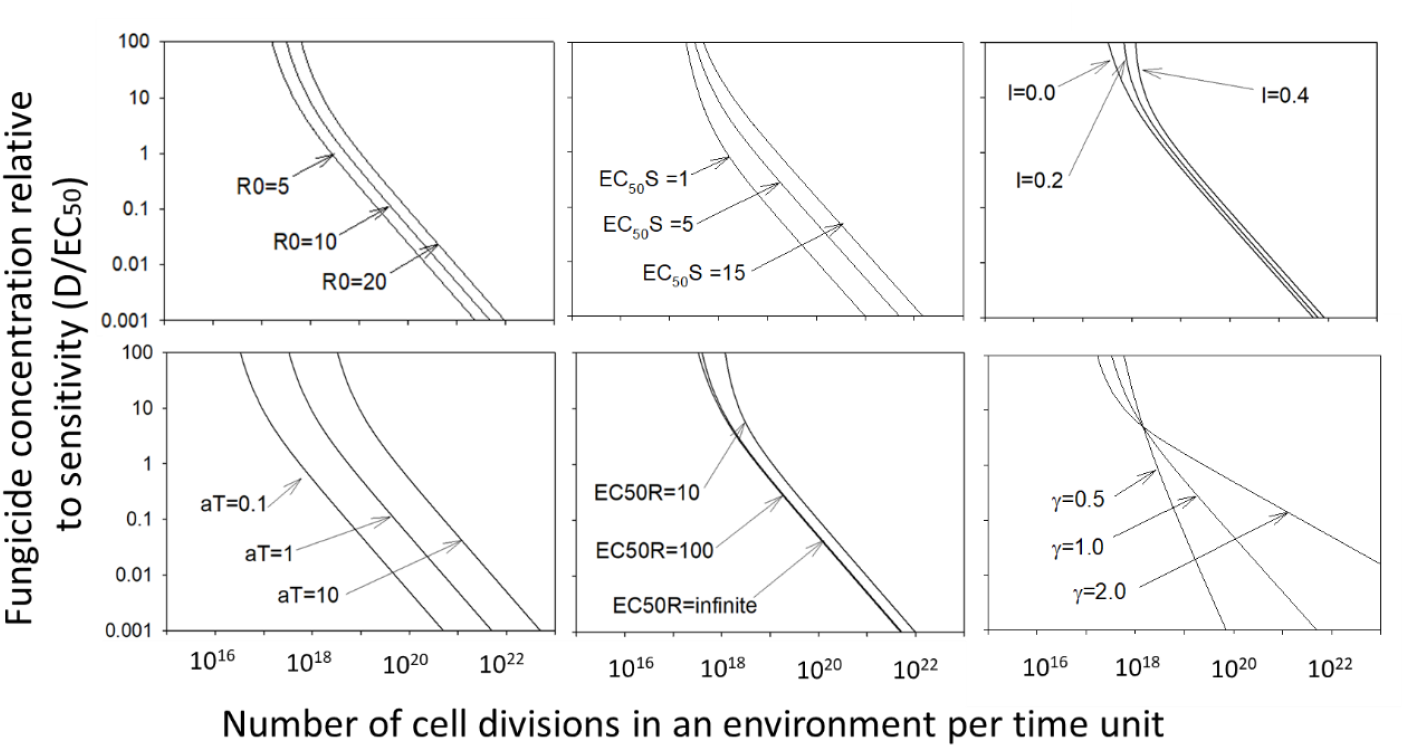
Effect on contours of relative emergence time of changing parameter values representing the net reproductive number (R_0_); EC_50_ of the sensitive strain; the asymptote of the dose response (I); The carbon absorption rate (aT); EC_50_ of the resistant strain (EC_50_R); and the dose-response curve shape parameter (*γ*).

Little is presently known about fitness costs to resistance in *Aspergillus* (Rivelli Zea et al., 2022; Chen et al., 2023), though fitness costs have been measured for azole-resistant *Aspergillus fumigatus* strains (Chen et al., 2023) and are known for plant pathogens quite widely. We therefore need to consider the possible effects of fitness costs. To this end we introduced a simple way of including fitness costs. When in the future more is known about fitness costs these can be considered in the model as described. A fitness cost was introduced as a reduction in the basic reproduction number of the resistant strain as R0R=f R0S. Including a fitness cost for the resistant strain of 5 % (f=0.95) substantially altered the emergence time contours (Figure 7). Even such a small fitness cost resulted in an infinite emergence time for fungicide concentrations below approximately 0.1 (D/EC50), because R0R is lower than R0S despite the presence of a low concentration of fungicide. Chen et al. (2023) show that such fitness cost values are possibly realistic for the azole-resistant *Aspergillus* case.

**Figure 7.**
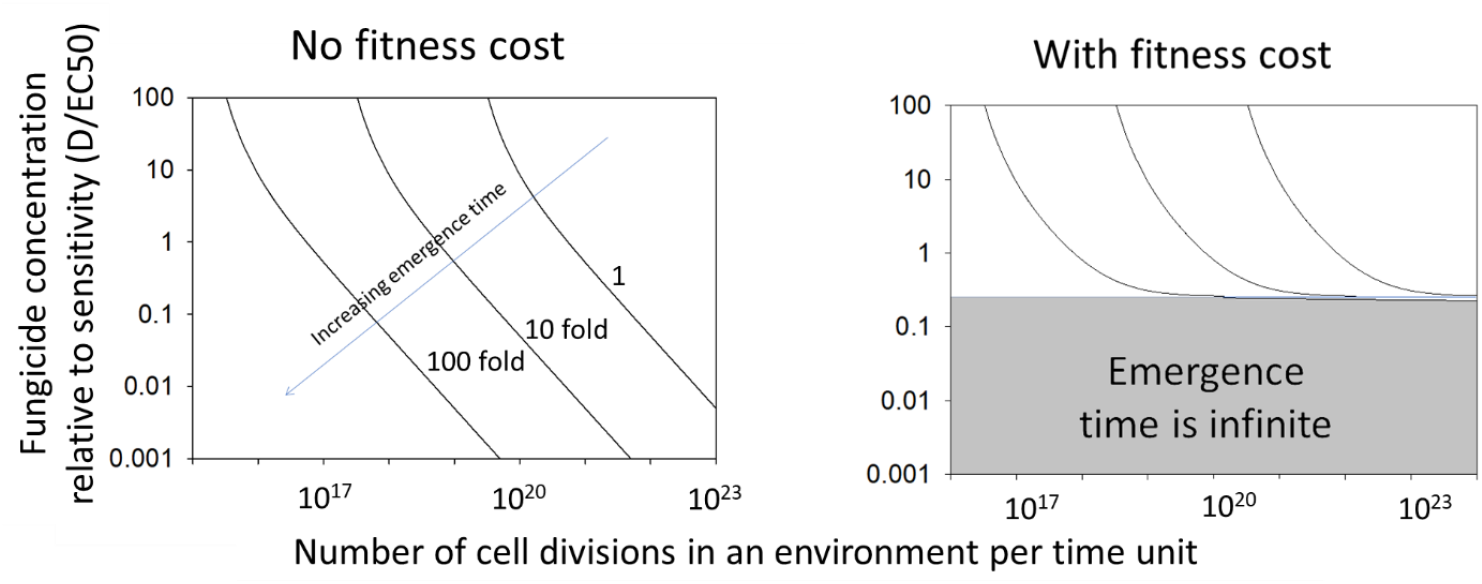
Contrast between emergence time contours without and with a small fitness cost for the resistant strain. Emergence time is infinite in the grey shaded area. The fitness cost is 5%, see main text for explanation.

#### Selection rate for fungicide resistance

Selection rate is independent of the size of the pathogen population and number of cell divisions, but sensitive to fungicide concentration; hence the horizontal contours for selection rate (S) in Figure 8 left. A small fitness cost is sufficient to stop selection at low fungicide concentrations (Figure 8 right).

**Figure 8.**
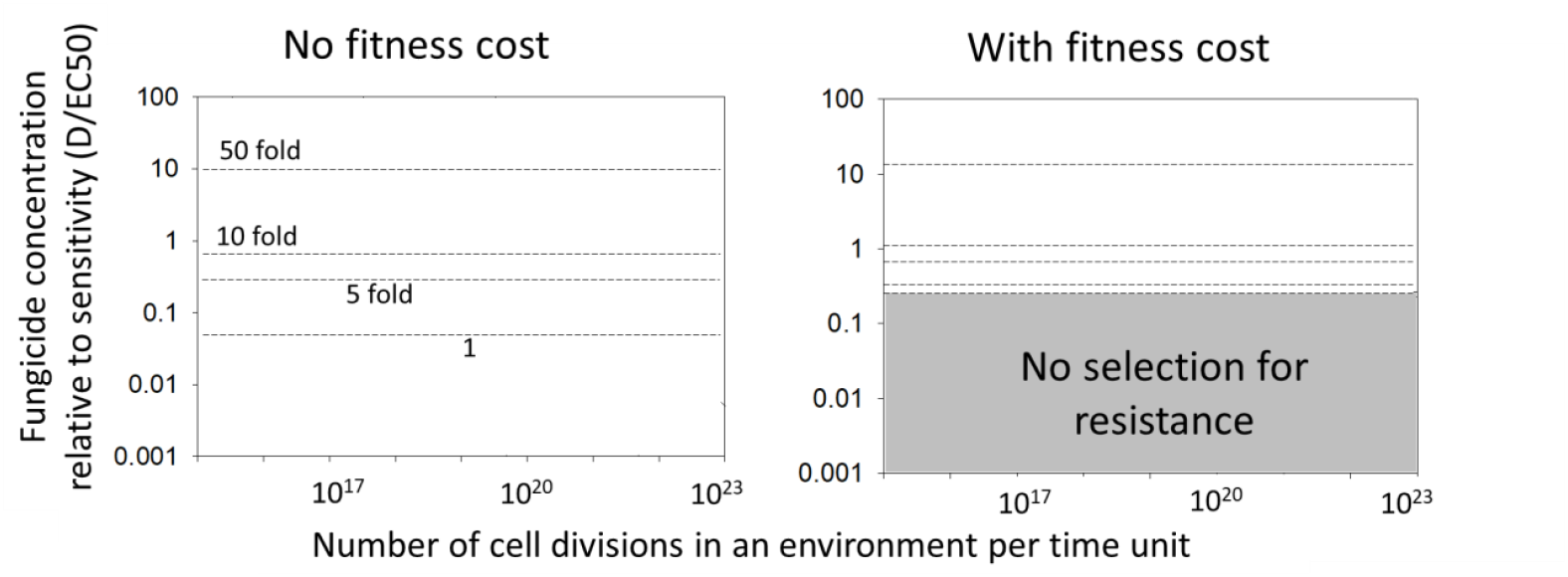
Selection rate (S) contours, with and without fitness cost of resistance. The fitness cost was 5%, see main text for explanation.

Sensitivity to parameters was explored by plotting (Figure 9) selection rate on fungicide dose (expressed relative to the sensitivity of the sensitive strain; EC_50_S). Selection rate was faster with increasing EC_50_ of the resistant strain and with higher shape parameter (*γ*) values for the dose response, which result in greater efficacy at a given dose. Selection rate was slower with increasing carbon absorption rate (*aT*) and with larger asymptote (*L*) values for the dose response, which result in lower efficacy at a given dose. If there is no fitness cost, the selection rate is independent of R_0_.

**Figure 9.**
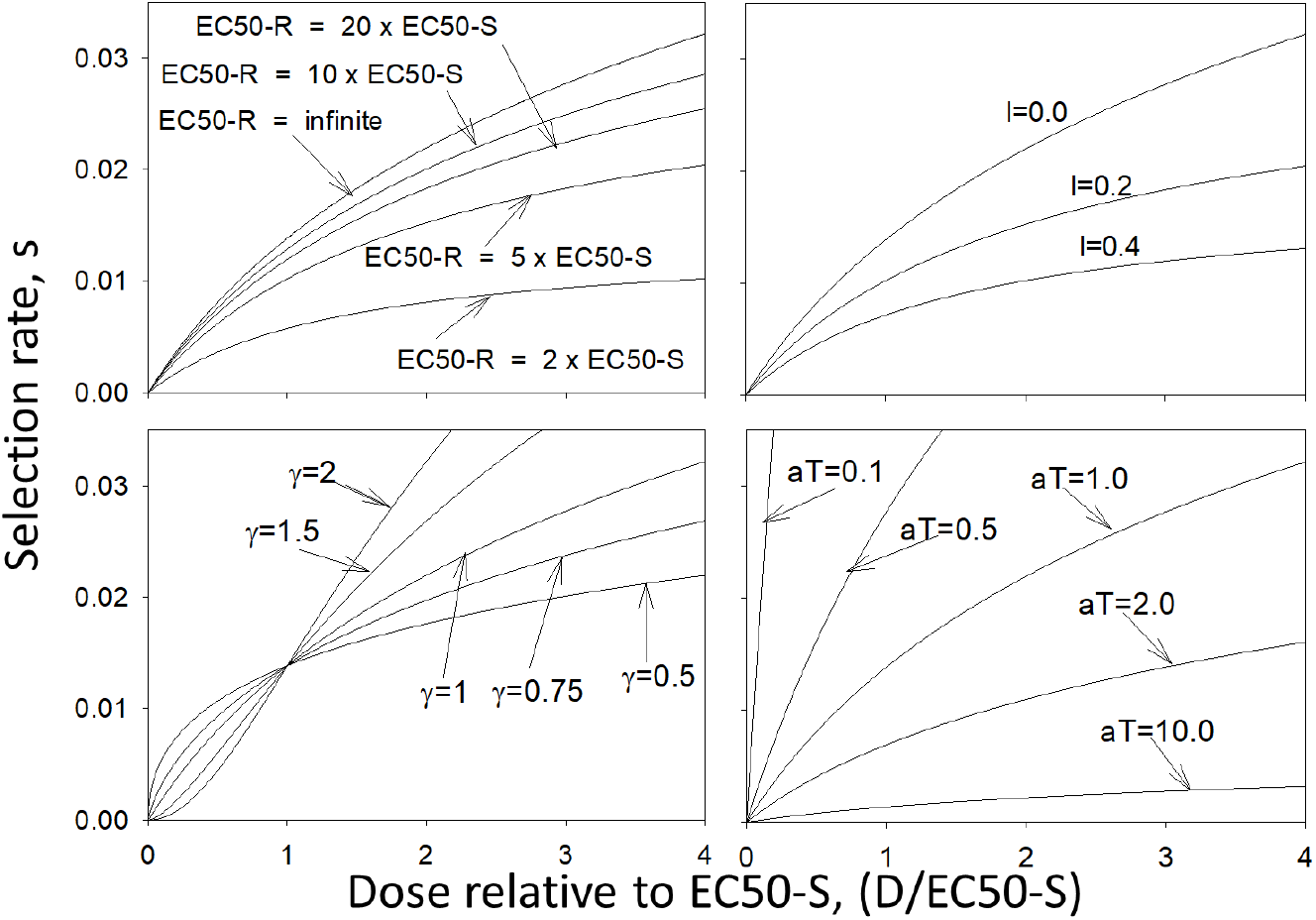
Relationships of selection rate on dose (relative to the EC50 of the sensitive strain) across a range of parameter values.

### Observed values for fungicide concentration and pathogen reproduction: example case for *A. fumigatus* in the Netherlands

This section compares landscape environments for pathogen population reproduction and fungicide concentration, as key determinants of emergence time and selection rate. Data for pathogen abundance (CFU/gr substrate) were found for *A. fumigatus* for many substrates across several countries. Values and literature sources are given in Supporting Information S3 and the ranges for key environmental compartments are summarized in Table 4 below. The values from different studies differ hugely, leading to the large ranges of values in Table 4. Nevertheless, the abundance in composts (from various types of plant material, including bulb waste) is orders of magnitude greater than other environments.

**Table 4.** Summary ranges of *A. fumigatus* abundance (colony forming units, CFU, per gram of substrate) in environments. Data for crop plants is in brackets, as there is no published evidence that reproduction is occurring on leaves.

| Environment | Pathogen abundance, CFU/gr |
| --- | --- |
| Compost (incl. flower bulb waste) | 40,000 to 800,000 |
| Agricultural field soil | 4 to 321 |
| Forest, meadow, grassland soil | 1 to 500 |
| [Crop plants on leaves] | [9 to 3,000] |

As discussed, emergence is sensitive to the total amount of reproduction (the total number of cell divisions in an environment per time unit). We thus need to convert the CFU per gram and the size of the environment (number of grams in the entire environment) into a rate of reproduction. In the supporting information we show how the total number of cell divisions in an environment is proportional to the CFU in the environment. We use this to calculate the scaled number of cell divisions in an environment; the calculation is summarized here.

With each cell division there is a small probability, ε, that a mutation (or other genetic event) occurs that makes the daughter cell resistance/less sensitive to the fungicide. The number of cell divisions in the fungal population is proportional to (i) the fungal biomass (per gr substrate), *B*, and (ii) the rate of reproduction per fungal individual/cell, *b*. Multiplying with the total number of grams of substrate in the environment under consideration, *A*, we find the total number of cell divisions in that environment. So

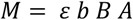

The per capita growth rate, *r*, of the fungus is the birth rate, *b*, minus the death rate, *μ*. Using the equations derived in the appendix we have

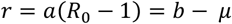

So *b = a*(*R*_*0*_ − 1) + *μ* and this equals 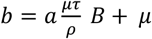 substituting in the above gives

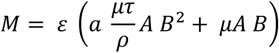

and since *B* is large we have

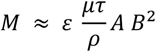

*B* is proportional to CFU as *B*=*ψ* CFU so we finally get

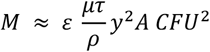

We will loosely term *A CFU*^*2*^ the ‘scaled number of cell divisions in the environment per time unit’. This is plotted on the x-axis of figure 10.

**Figure 10.**
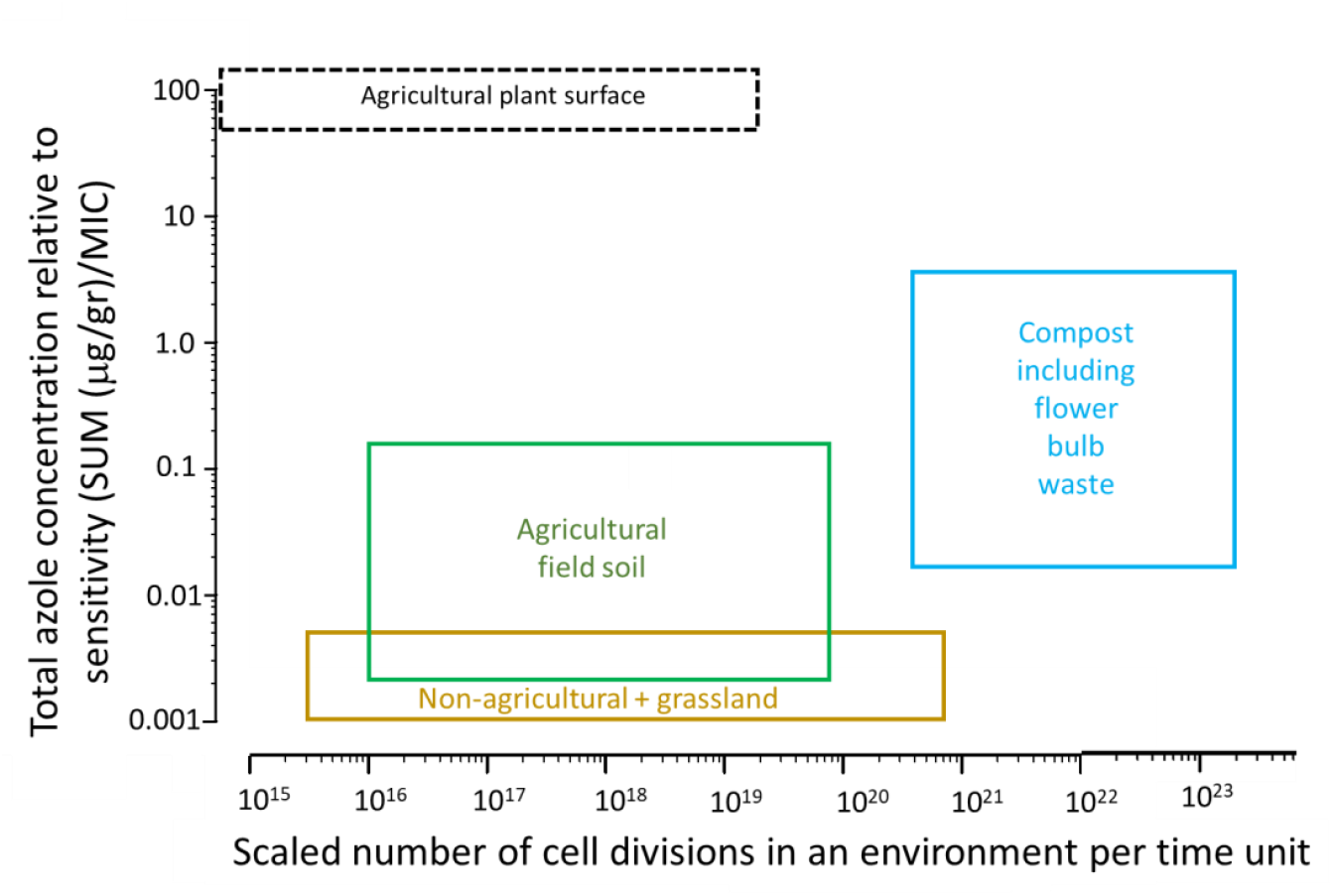
Example case for *A. fumigatus* for the Netherlands, illustrating the key risk variables for four environmental compartments. The box for each environment represents the wide range of values from the literature for fungicide concentration and pathogen reproduction. Any individual case should be somewhere within the relevant box. The agricultural plant surface environment is shown within a dotted line, as *A. fumigatus* is not known to reproduce in that environment, in which case that environment could not drive resistance emergence or selection.

Data needed to estimate the weight of substrate (e.g. the total amount of compost in a country) to calculate the relative population rate of reproduction (CFU per unit time). A search for such data across multiple countries found suitable data for the Netherlands (see Supporting information S3). Values were available for weight of compost where the source material was from agricultural fields, which were likely to have been treated with fungicide. For other countries, data were either not found or were not feasible to categorise by specific environmental compartment with sufficient certainty. The results should therefore be treated with caution, as The Netherlands is unlikely to be representative of other countries (see discussion).

Literature sources and data on concentrations of agricultural azoles in multiple substrates are summarised in Supporting Information S4. To assess risk, fungicide concentration needs to be scaled by pathogen sensitivity, so data for sensitivity of *A. fumigatus* to agricultural azoles (Table 5) was obtained from Allizond et al., 2021; Cao et al., 2020; Fraajie et al., 2020; Garcia-Rubio et al., 2021; Kang et al., 2022; Jørgensen et al., 2021; Meneau & Sanglard, 2005; Prigitano et al., 2022; Roberts et al., 2020; Snelders et al., 2012.

**Table 5.** MIC and EC50 values for *A. fumigatus* and agricultural azoles. Sources cited in text. Values are either means of a range of values or a single value given in the source.

| Active substance | MIC | EC<sub>50</sub> |
| --- | --- | --- |
| Prothioconazole | 7.13 | 0.29 |
| Prothio-desthioconazole | 0.5 | Not reported |
| Prochloraz | 0.30 | Not reported |
| Propiconazole | 3.22 | 5.0 |
| Tebuconazole | 2.67 | 4.5 |
| Cyproconazole | 32 | Not reported |
| Flusilazole | 2.1 | Not reported |
| Difenoconazole | 1.47 | 2.5 |
| Epoxiconazole | 3.42 | 9.0 |

Summing concentration data for all azoles from Supporting Information S4 and dividing by MIC values in Table 5, gave the indicators of azole efficacy in Table 6.

**Table 6.** Sum over all azoles (ug/gr)/MIC as an indicator of azole efficacy. The full list of values and literature sources are given in Supporting Information S3. Data from crop plant leaves is in brackets as the environment may not be relevant, as *A. fumigatus* is not known to reproduce in that environment.

| Environment | $\Sigma(\text{ug/gr})/\text{MIC}$ |
| --- | --- |
| Compost (incl flower bulb waste) | 0.024 - 6.6 |
| Agricultural field soil | 0.0022 - 0.20 |
| Forest, meadow, grassland soil | 0.001 - 0.004 |
| [Crop plants on leaves] | [20.3 - 34.1] |

The ranges of values for rate of reproduction and total azole concentration, scaled by MIC, for the four landscape environments are plotted in Figure 10. The ranges of values from the literature are huge (noting the log scale of both axes). It seems plausible that a true value from any specific substrate and location should fall somewhere within the relevant box. As shown previously in Figures 7 and 8, the top right of the chart represents a region of high risk (short emergence time and fast selection) and the bottom left represents a region of low risk (long emergence time and slow selection). If there is a fitness cost for a particular resistant strain, then the lower part of the chart may represent a region of no risk from that strain.

Compost (including flower bulb waste) is positioned towards the high risk end of the spectrum, where short emergence times and fast selection may be expected. Leaves of crop plants are also, in principle, a high risk environment, due to the high concentration of fungicide directly applied to the crop canopy. However, there is no convincing evidence in the literature that *A. fumigatus* reproduces in that environment, making emergence or selection highly unlikely. Despite the large ranges of values for the environments, compost is separated from ‘agricultural field soil’ and ‘non-agricultural land and grassland’; which are both positioned more towards the lower risk end of the spectrum, due primarily to low fungicide concentrations. However, the risk from these environments cannot be entirely discounted – there is a spectrum of risk. Pathogen abundance values per unit of substrate in soil and non-agricultural/grassland environments are orders of magnitude lower than for compost, but the amount of substrate is far greater than compost as the land areas occupied are large.

If resistant strains typically carry even a small fitness cost, then Figure 10 would need to be interpreted substantially differently. Soils, non-agricultural environments and grassland may fall in the region of no risk, due to low fungicide concentrations; making compost the sole hotspot in the Netherlands.

## DISCUSSION

The resistance risk posed to medicine from agricultural use of fungicides depends on the traits of the specific human pathogen, the fungicide and the landscape environments in which they occur. Combinations of traits which result in high or low risk have previously been termed ‘hotspots’ or ‘coldspots’, respectively (Gisi, 2022). A combination of traits can make resistance emergence and selection, and exposure of humans, sufficiently unlikely that a pathogen or pathogen/fungicide/environment combination can be considered a coldspot. Current evidence suggests that seven species (six *Candida* species and *Pneumocystis jirovecii*) of the 27 human pathogens considered here, each lack at least one of the mechanisms necessary for them to pose a resistance risk to humans from agricultural fungicide use. However, the degree of evidence required to prove the absence of a mechanism, and hence the absence of risk, is difficult to define. This is a particular concern for pathogens, such as *Candida auris*, which are ranked as a critical risk to human health (WHO, 2022).

For the other 20 human pathogens, there is limited evidence for trait combinations which identify the potential for some degree of risk. Hotspots and coldspots represent the ends of a spectrum of risk. The quantitative analysis presented here develops an approach to identify the traits of importance in determining the degree of resistance risk. The analysis method could be applied across a range of pathogens and environments. Before considering the findings, and their implications for risk management, we first consider limitations of the analysis.

A first major constraint on risk analysis is that there are very substantial evidence gaps about the behaviour of human pathogens in landscape environments, as this was only recently recognised as a relevant area of study. Some of these gaps are qualitative, for example, it would be valuable to corroborate the hypothesis that the surfaces of agricultural plants do not provide a suitable environment for reproduction of *A. fumigatus* or other human pathogens. This is currently based on the low density of *A. fumigatus* found on leaves (Supporting Information S3) and on what is known about the conditions required for reproduction. Crop canopies in agricultural fields are a very large environment and have higher fungicide concentrations than other landscape environments, so the presence or absence of pathogen reproduction (as opposed to the presence or absence of the pathogen) makes a critical difference to the analysis of the risk of development of resistance due to agricultural use of fungicides. Other knowledge gaps are quantitative, both about the characteristics of the specific pathogens and landscape environments and, critically, about the flows of substrate, pathogen and fungicide which link environmental compartments.

These gaps limit parameter estimation. Plausible parameter ranges for sensitivity analysis were therefore obtained for a taxonomically diverse range of fungi (mainly crop pathogens) which have been widely studied in the field. The parameter range for human pathogens may differ. The model developed and the analysis were constrained to single, rather than coupled, environments. In reality, the flow of substrate, pathogen and fungicide between environmental compartments may materially affect the degree of resistance risk. Certain environments are likely to be more or less conducive to emergence and selection and other environments may have a larger contribution to human exposure, so different phases may occur sequentially in different environments; linked by transfer between environments.

A second constraint on risk analysis is the stochastic nature of resistance emergence. Whereas the selection phase is largely deterministic, the emergence phase is driven by random mutation events and the probability of progeny surviving from a mutation event (Hobbelen et al., 2014; <u>Mikaberidze</u> et al., 2017). For any given set of conditions, emergence may happen substantially more or less quickly by random chance. The lack of experimental studies on emergence in relevant environments means that insights are dependent on unvalidated modelling studies. We should, therefore, be cautious about using the current findings to guide practical decisions and recognise that stochasticity will always limit the extent to which we will be able to predict resistance outcomes.

However, the current case of parallel development of DHODH antifungals and fungicides (Verweij et al., 2022) illustrates that there is an urgent need to support decisions on whether it is safe to develop fungicides for agricultural use from MoA which are also being developed for pharmaceutical use. There is also a pressing need to identify effective and practical methods to mitigate risk. What can be taken from this preliminary quantitative analysis to support such decisions and focus future work?

The risk of resistance emergence was found to be strongly affected by the amount of pathogen reproduction in an environment, fungicide concentration relative to the sensitivity of the pathogen and fitness cost. Emergence was moderately affected by the carbon absorption rate and to the shape of the dose response curve when fungicide concentrations were low. The implications of each of these are considered below.

### Amount of pathogen reproduction in an environment

Emergence is determined by the number of resistance mutations and the probability of mutant progeny surviving. The main sub-components being, the size of the environmental compartment, the density/abundance of the pathogen in the environment, the mutation rate and R_0_. Previous work on resistance risk has focussed on the density of the pathogen in the environment, but there is now an open question about the relative resistance emergence risks posed by a small environment with a high pathogen density versus a large environment with a low pathogen density. Data on the size of environments and the density of human pathogens within each environment would be valuable to assess relative risks and hence focus mitigation. For the example case of *A. fumigatus* for the Netherlands, it appears that the very high pathogen density in plant waste can outweigh the small size of the environment. The Netherlands is atypical in the intensity of horticultural production, and flower bulb production in particular. Data from countries with differing cropping would aid risk assessment.

In principle, a doubling of mutation rate would halve the predicted emergence time. Although mutation rates can be quantified in general, across the genome (e.g. Bridges, 1975; Amaradasa & Everhart, 2016), quantification of mutation rates in fungicide target sites is lacking. Nevertheless, there is concern about the effects of ‘sub-lethal’ doses of fungicides on mutations. Mutagenesis is a red line for pesticide regulators, but it is argued that fungicides can act as a source of stress, resulting in a higher mutation rate (Gressel, 2011; Verweij et al., 2016; Amaradasa & Everhart, 2016; Gambhir et al., 2021).

### Fungicide concentration relative to sensitivity

The concentration of fungicides in different environments has already been recognised as a key risk factor (Gisi, 2022), but the relationship between concentration and resistance risk is poorly understood. Most of the work on the effect of fungicide dose in field environments has been to understand fungicide efficacy against crop pathogens, within the range of doses typically applied to crops. *In vitro* studies have quantified sensitivity phenotypes of wild type and resistant strains; usually by measuring growth across a log scale of doses, fitting a log-logistic dose response curve and summarising sensitivity by a metric derived from the curve, such as MIC (Fraaije et al., 2020). Relating *in vitro* sensitivity to effects in the field has proved uncertain, although such relationships have been found for crop pathogens (Blake et al., 2017). In landscape environments other than crops and compost/bulb heaps, doses are so small that it is probably impossible to measuring their effects on human pathogens in the field.

Nevertheless, the analysis presented here suggests that, in the absence of fitness costs, low concentrations can still affect the risk of emergence. These concentrations are distant from the EC_50_ or MIC values typically used to summarise sensitivity. A ‘no effect concentration’ (NOEC or NEC) on the wild type or ‘predicted no effect concentration’ (PNEC) on the wild type may be more relevant metrics. Fungicide sensitivity metrics are considered further, in relation to resistance risk, in Supporting information S6.

### Shape of the dose response curve

The finding that emergence time is moderately affected by the shape parameter of the dose response relates to the preceding section on fungicide concentration relative to sensitivity. In the model, concentration was scaled by fungicide sensitivity, with the latter quantified by EC_50_. The effect of a given dose on growth will vary depending on the shape parameter, particularly at doses distant from the inflection point of the response curve at which EC_50_ is defined – such as the low doses found in landscape environments.

### Fitness cost

The effect of fitness cost interacts with fungicide concentration such that at low concentrations the emergence time can become infinite, because the sensitive strain is more fit than the resistant strain at very low concentrations. The fitness costs reported for *A. fumigatus* (Arendrup et al., 2010; Chen et al., 2023) seem large enough to make it possible that emergence could not have occurred in landscape environments with low fungicide concentrations, such as agricultural soils and semi-natural environments. Other authors (Gonzalez-Jiminez et al 2021) have inferred that rapid dispersal of azole resistant *A. fumigatus* strains supports the hypothesis that there is no diminution in fitness.

Understanding the effect of fitness cost may help to explain in retrospect what has been seen but exploiting it to guide decisions about development of new MoAs will be challenging. Such decisions would be based on information that can be obtained during the development of new active substances, rather than being a *post-hoc* assessment after product release. Laboratory mutation studies can often generate a range of resistant mutants. Fitness studies on these may provide some useful insight, although there are many cases for crop pathogens where target site mutations have occurred in the field, that were not found by laboratory studies (Hawkins & Fraaije, 2016).

### Carbon absorption rate

Too little is known to assess the practical relevance of this parameter for human pathogens in landscape environments.

Selection for a resistant strain is not determined by the size of the environmental compartment, nor by the density of pathogen in the environment. Risk of selection was found to be strongly affected by fungicide concentration relative to the sensitivity of the pathogen and to fitness cost. Selection was moderately affected by the ratio of fungicide sensitivity of the sensitive and resistant strains, the asymptote of the dose-response curve, the shape parameter of the dose-response curve and the carbon absorption rate.

### Fungicide concentration relative to sensitivity

Fungicide concentration and sensitivity combine to affect the *per capita* growth rates of the sensitive, r_S_, and resistant, r_R_, strains. If growth of the resistant strain is almost unaffected at field doses (i.e. a high resistance factor), then the higher the concentration and the greater the sensitivity of the sensitive strain, the greater the difference between r_S_ and r_R_, and the faster the rate of selection. The converse will be true at low concentrations. This logic explains other findings from the analysis. Firstly, the rate of selection was also found to be moderately affected by the ratio of fungicide sensitivity of the sensitive and resistant strains. If, for example, the resistance factor is low, then r_R_ at a given dose will be lower and the difference between r_S_ and r_R_ will be smaller. Secondly, the rate of selection was found to be moderately affected by the asymptote and shape of the dose response, as (for a given EC_50_ value) these parameters relate to the sensitivity of the pathogen at a given dose – particularly at doses distant from the EC_50_.

### Fitness cost

The effect of fitness cost interacts with fungicide concentration such that at low concentrations, the rate of resistance selection can become nil (or the wild-type may outcompete the resistant strain). In the absence of fungicide the resistant population, carrying a fitness cost, will grow more slowly than the wild-type population. At high fungicide concentrations the resistant strain will grow and reproduce more quickly, outcompeting the wild-type (mutants with large or lethal fitness costs are will not emerge or be selected for). The fitness advantage of the resistant strain in the presence of fungicide will diminish with dose until at a certain dose the fitness of the two strains will be the same. Below that dose, the fitness of the wild-type will exceed that of the mutant. The fitness costs reported for *A. fumigatus* (Arendrup et al., 2010; Chen et al., 2023) seem large enough to make it possible that selection could not have occurred in landscape environments with low fungicide concentrations, such as agricultural soils and semi-natural environments. This fits with experimental and observational evidence that increases in azole resistance in *A. fumigatus* could not be detected in soil in the presence or absence of fungicide treatment (Barber et al., 2020; Fraaije et al., 2020; Godeau et al., 2023 and other literature cited therein), though Barber et al. (2020) measured a slight, transient increase in the proportion of ARAf in one of two seasons.

The scope of our analysis was limited to landscape environments, where the output variable is the emergence and rate of increase of the resistant fraction in the pathogen population. Given a mechanism for transfer from the landscape to humans, higher resistant fractions result in higher risk of resistant fungal infections. Understanding the coupling between the landscape and medical settings will be critical, to be able to quantify the flow of pathogen propagules and its consequences for the probability that a human is infected with a resistant pathogen strain. The size of the pathogen population in the landscape is not relevant to selection but will be relevant to human exposure.

Generic principles or guidance can probably be defined for assessing the safety of dual use, but assessments will need to be made separately for each combination of human pathogen, agricultural fungicide active substance and medical anti-fungal active substance. This will require data to be gathered to agreed protocols and shared at an early stage in product development. The data required may differ according to the life cycle traits of the pathogen under consideration.

The analysis broadly supports the conclusions of other workers (Gisi et al., 2022; Verweij et al., 2022) about the factors to be considered in risk assessment. Decisions on the safety of dual use should, as a minimum, account for the following components: (i) presence/absence of cross-resistance between the agricultural fungicide and medical anti-fungal active substances, (ii) importance of the anti-fungal for control of the human pathogen, (iii) importance of the fungicide for sustainable control of crop pathogens, (iv) size of the human pathogen population in each relevant landscape environment (size of environment and density of pathogen), (v) presence/absence of human pathogen reproduction in that environment, (vi) the amount of the fungicide in each environment, relative to the sensitivity of the wild-type human pathogen, (vii) dispersal of the pathogen from an environmental compartment to create human exposure, and (viii) the feasibility of mitigation measures.

Given the qualitative and quantitative knowledge gaps, achieving consensus about acceptable safety limits will be challenging. For example, the analysis suggests that (in the absence of fitness costs) doses substantially below the MIC could cause significant selection in environmental compartments (given that the pathogen reproduces in that compartment) with considerable reproduction of the human pathogen. Doses in an environmental compartment would need to be below the chronic no effect concentration for the wild-type human pathogen, for selection to be negligible.

A separation of MoA used in medicine and agriculture would provide a complete solution to the medical risk, but if it is achieved by preventing MoA being developed for crop protection, then it creates risks to food production. Balancing these risks is a social and regulatory problem.

Abandoning the agricultural use of a MoA to which resistance has already developed in a human pathogen is likely to have limited benefit as a mitigation measure. It may prevent a further increase in the resistant fraction, but unless there is a substantial fitness cost associated with resistance, the frequency of the resistant strain would only decline slowly in the landscape. If there is a substantial fitness cost, then most landscape environments (with low fungicide concentrations) are unlikely to be the primary source of the problem.

Doughty et al. (2021) reviewed risk mitigation for azole resistant *A. fumigatus*, focusing on plant waste ‘hotspots’ that are particularly conducive to high densities of that pathogen. More broadly, across pathogen species and MoA, our analysis suggests that mitigation measures will be most effective if focused on: (i) reducing the amount of pathogen reproduction (through size of environment or pathogen density) - to slow emergence, (ii) reducing the concentration of fungicide in landscape environments that are conducive to pathogen reproduction - to slow emergence and selection, and (iii) developing active substances for agricultural use to which human pathogens have low sensitivity - to slow emergence and selection. How such mitigation might be implemented in practice will differ depending on the characteristics of particular pathogens. The analysis provides a method of exploring which environmental compartments pose a particular risk, but has highlighted the lack of information from landscape environments for human pathogens of concern.

The analysis presented here indicates the size of the reductions that would be necessary to have a substantial effect on the resistance risk for human pathogens from agricultural fungicide use. In many cases the effects are non-linear, so order of magnitude reductions would be needed to have a practically useful effect on risk. There are examples where such differences might be achieved. For example, active substances within the same MoA can differ by orders of magnitude in their effect on a given pathogen. Identifying specific high risk environments within the landscape may also hold significant potential for resistance risk mitigation.

## Acknowledgements

This work was funded by CROPLIFE INTERNATIONAL aisbl.

## Data availability statement

Data sharing is not applicable to this article as no new data were created or analysed in this study.

## Supporting Information legends

**Supporting Information S1** Evidence for a potential mechanistic route for resistance from agronomic settings to medical settings

**Supporting Information S2** Model derivation

**Supporting Information S3** Values and sources for pathogen density (CFU/gr) and number of pathogen cell divisions in different environments

**Supporting Information S4** Values and sources for azole concentrations in different environments and substrates

**Supporting Information S5** Values for the curvature parameter of the dose-response

**Supporting Information S6** fungicide sensitivity metrics

## Supporting Information S2

Model derivation

The equations for the rate of change of the densities shown in Figure 3 are:

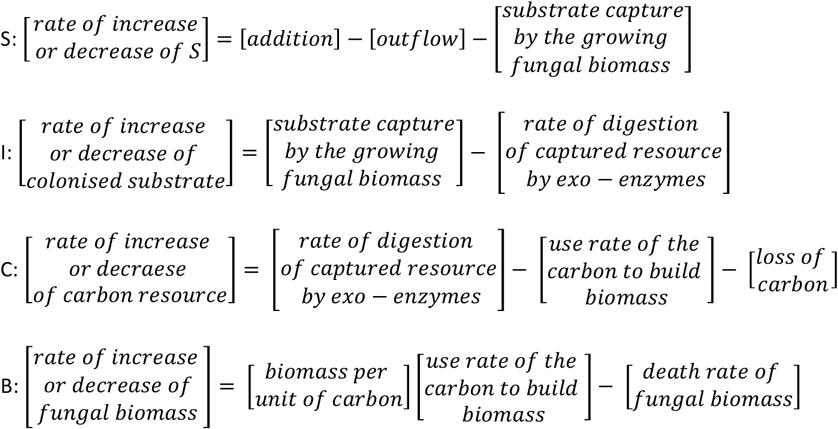

Assumptions:

The substrate addition rate is a constant θ, so

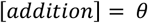

The *per capita* outflow rate is ρ, so

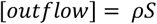

One unit of carbon used for fungal growth increases the size of the fungal mass by α. So,

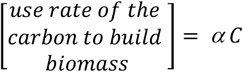

The amount of substrate colonised by one unit of biomass growth is τ. So,

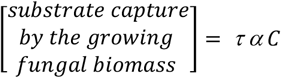

The digestion of substrate into carbon usable to the fungus is mediated by an exo-enzyme. Therefore, we model the rate of usable carbon formation with Michaelis-Menten dynamics.

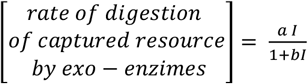

Carbon usable to the fungus can flow into the environment. It is also usable to other competing micro-organisms. The per capita rate of carbon loss is *ξ*, so

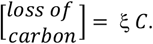

Each unit of fungal biomass has a probability *μ* per time unit to dy. So,

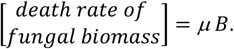

The model of linked differential equations thus has the form

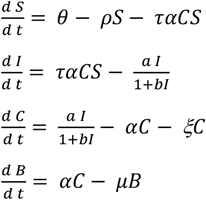

The net-reproductive number, *R*_0_, (the number of daughter lesions per mother lesion when availability of susceptible host is not limiting) is given by

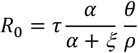

The exponential population growth rate, *r*, is

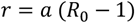

First note that in steady-state the density of S, *Ŝ*, is given by

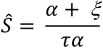

and

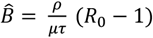

### Representing fungicide action

To represent fungicide effect, a log-logistic dose response curve is given by

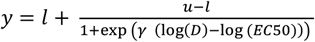

Which is the same as

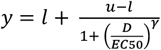

Where EC50 is the EC_50_, *D* is fungicide dose, *l* is the asymptote, *u* is the value of *y* at *D*=0 and *γ* is a shape parameter, determining the length of the plateaux around *D*=0. We will take *u*=0 so y=1 at *D*=0. *y* can then be interpreted as the relative response of the fungal density to fungicide dose *D*.

The response of fungal growth to fungicide is measured in dose response studies, in which a range of doses are applied and density is quantified after a time interval, resulting in a response curve. As dose response studies are usually conducted with a low starting density, we assume exponential fungal biomass growth over the period, *T*, from fungicide application to assessment of fungal density in a dose response experiment, hence

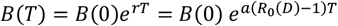

Then *R*_0_(*D*) is given by

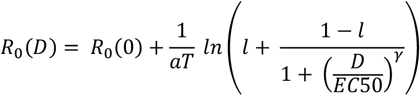

and

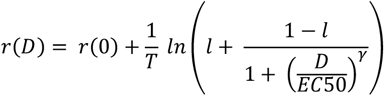

### Calculating emergence time for fungicide resistance

*P* is the probability that a resistant mutant develops a population of a size not prone to random extinction. *M* is the number of resistance mutations in the sensitive pathogen population per time unit. The mean time until resistance emerges, *T*_emerge_, is given by (<u>Mikaberidze</u> et al., 2017)

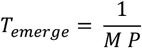

We derive *P* and *M* and finally *T*_emerge_.

#### Deriving P

Following Mikaberidze et al. (2017) the probability, *P*, that a mutant will emerge into a completely sensitive population is

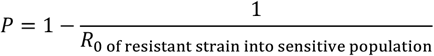

Where R_0 of resistant strain into sensitive population_ is the net-reproductive number of the emerging resistant strain in a population consisting of the sensitive pathogen only. Using the model this leads to

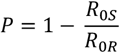

Where R_0S_ is the net-reproductive number of the sensitive strain (in the absence of the fungus), and R_0R_ the net-reproductive number of the resistant strain.

Now using our expression for *R*_0_ we get

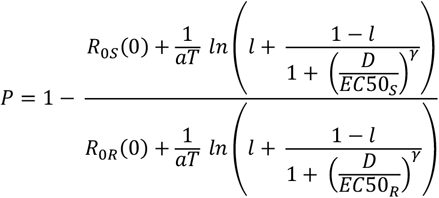

Where EC50_S_ and EC50_R_ are EC_50_ values of the sensitive and resistant strain, respectively. If there are no fitness costs, then *R*_0S_=*R*_0R_.

#### Deriving M

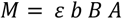

The per capita growth rate, *r*, of the fungus is the birth rate, *b*, minus the death rate, *μ*. Using the equation above we thus have

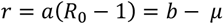

So *b = a*(*R*_*0*_ − 1) + *μ* and this equals 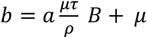 substituting in the above gives

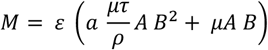

And since B is large we have

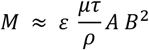

B is proportional to CFU as B=Ψ CFU so we finally get

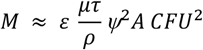

We will loosely term *A CFU*^*2*^ the ‘scaled number of cell divisions in the environment per time unit’.

#### Calculating T_emerge_

An analytical solution can be derived for emergence time (avoiding the need for simulation) thus

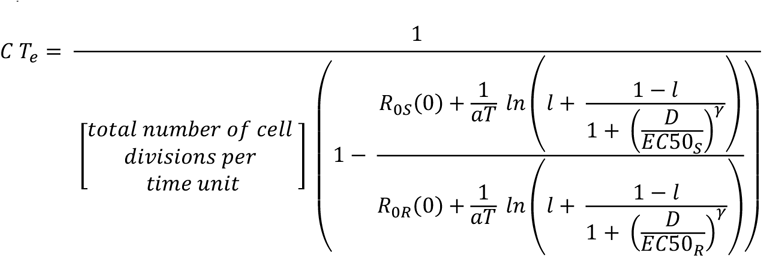

where 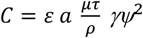.

The emergence time *T*_*e*_ has a constant in front of it which is not known, so we can only plot lines of equal emergence time multiplied with an unknown constant.

### Calculating rate of selection for fungicide resistance

Selection is measured by *s = r*_*R*_ *− r*_*S*_ (Milgroom and Fry, 1988). For the model of a fungus in a given environment we get

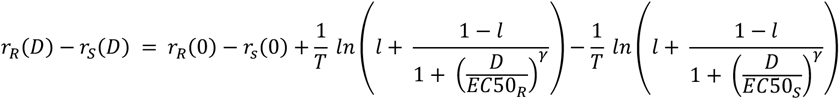

When there is no fitness cost to being resistant *r*_R_=*r*_S_ in the absence of fungicide. If there is no fitness cost and the dose response curve has asymptote at zero and the resistance is absolute 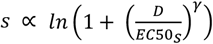

## Supporting Information S3

Values and sources for pathogen density (CFU/gr) and number of pathogen cell divisions in different environments

*A.fumigatus* densities in a range of environments, with source literature, are given in the table below.

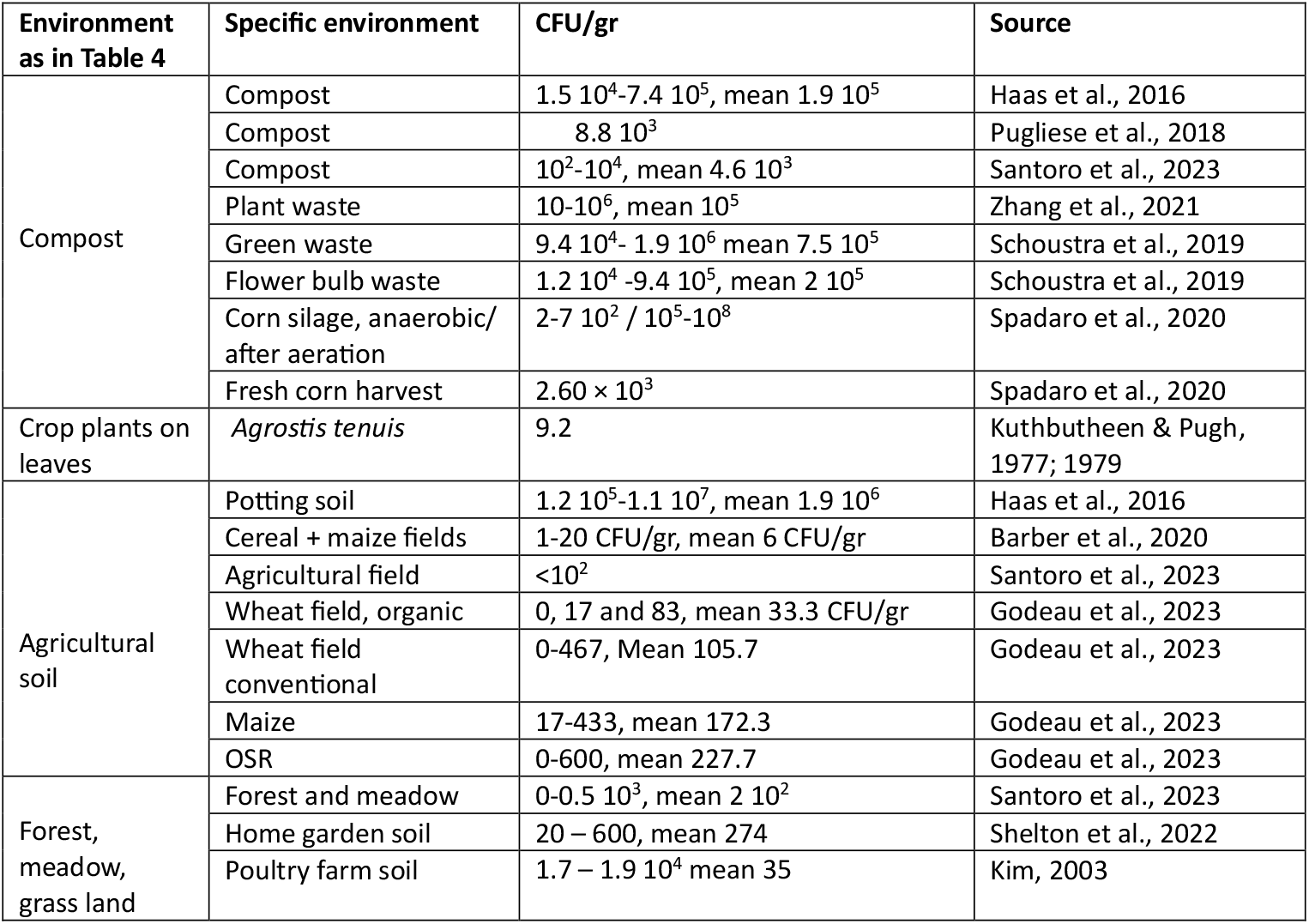

### Calculation of the weight of each specific environment for The Netherlands

#### For soil

land area of the Netherlands: 33,500 km^2^; Agricultural area of arable crops: 5,290 km^2^; Built-up area: 5,192.5 km^2^ (Wikipedia). Non-agricultural area (semi-natural + grassland): 22,887.5 km^2^

The top 10 cm depth was taken to be most biologically active and likely to release propagules through cultivation. Weight by volume data for soils were used to calculate total weight of soil for each environment.

### For compost

Calculation 1:

Bulb waste:

Roelofs & Gude (2013): Flower bulb waste is 74,200 tonne dry matter. • Franssen

(1999) ratio dry weight: fresh weight bulbs is 0.4.

Gives 185,000 tonne bulb waste.

Vegetable waste:

Anonymous (2009): Vegetable waste is 216,000 tonne.

What is composted:

All flower bulb waste is composted. We have not found data on the use of bulb waste for biogas for the time period covered by the data. However, a fraction of the bulb waste is currently used for biogas production. WE note here that even when 50% of the bulb waste is used for biogas, the environmental compartment box for compost does not shift the left in figure 10 to an extent that our conclusions change.

Elbersen & Groen (2020), de Bruin & Marinissen (2020) estimate 48% of all

agricultural crop waste is composted. We use this for the vegetable waste.

Composted substrate per year is 185,000 + 0.48 x 216,000 = 288,680 tonnes

Calculation 2:

Total organic crop waste streams:

Meeusen-van Onna et al. (1998) total organic waste staying on farm is 342,000

tonne. From the waste leaving the farm 315,000 tonne goes to compost + fertiliser +

soil cover.

In total 657,000 tonne crop waste per year goes to compost + fertiliser + soil cover.

What is composted:

Elbersen & Groen (2020), de Bruin & Marinissen (2020) estimate 48% of all

agricultural crop waste is composted. We use this for the vegetable waste.

657,000 tonne x 0.48 = <u>3</u>15,360 tonne is composted.

Conclusion: 2.9 10^8^ – 3.2 10^8^ kg compost

*For crop leaves*:

Ha grown in the Netherlands: Total cereals: 161,206 ha

Maize: 13.742 + 183.274 + 6.582 = 203,598

Potato: 163.058 Leaf fresh weight:

Wheat: 9.5 10^3^ kg/ha. (Berghuis et al 2023)

Maize. 2 10^3^ – 2.5 10^3^ kg/ha (de Beus 2015, Plenet et al 2000)

Potato: 3.8 10^3^ – 5.0 10^3^ kg/ha (Ten Den et al 2022) Total weights:

Cereals: 1.2 10^9^ kg

Maize: 4.1 10^8^ – 5.1 10^8^ kg

Potato: 6.2 10^8^ – 8.2 10^8^ kg

Total: 2.23 10^9^ – 2.6 10^9^ kgWe used leaves here only because the references assessed Aspergillus on plant leaves. It is very, likely that Aspergillus will also be present on stems and other plant parts. We have left them out of consideration here.

### Combining the CFU/gr and the gr per environment to estimate reproduction

| environment | CFU/gr | gr in NL | (CFU/gr)<sup>2</sup> x gr in NL |
| --- | --- | --- | --- |
| Compost, including flower bulb waste | 3.8 10<sup>4</sup>-7.9 10<sup>5</sup> | 2.9 10<sup>11</sup> – 3.2 10<sup>11</sup> | 4.1 10<sup>20</sup> – 2.0 10<sup>23</sup> |
| Agricultural crop fields | 3.6 – 320.6 | 8.1 10<sup>14</sup> | 1.1 10<sup>16</sup> – 8.1 10<sup>19</sup> |
| Semi-natural and grassland | 1 -5 10<sup>2</sup> | 3.4 10<sup>15</sup> | 3.4 10<sup>15</sup> – 8.5 10<sup>20</sup> |
| Crop leaves | 9.2 – 2.9 10<sup>3</sup> | 2.2 10<sup>12</sup> – 2.6 10<sup>12</sup> | 1.9 10<sup>14</sup> – 2.2 10<sup>19</sup> |

## Supporting information S4

Values and sources for azole concentrations in different environments and substrates

Azole concentrations in a range of environments and substrates, with source literature, are given in the table below.

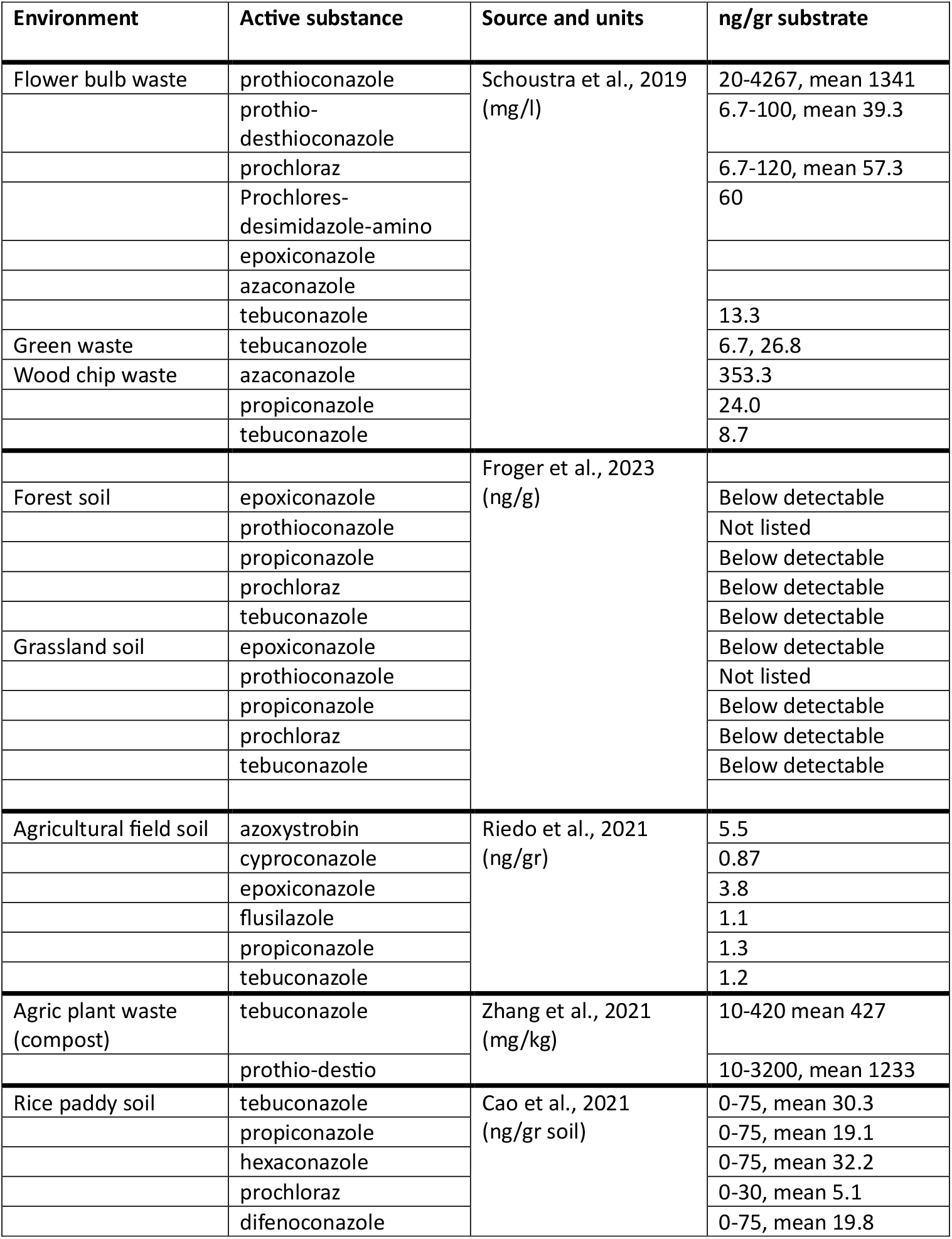
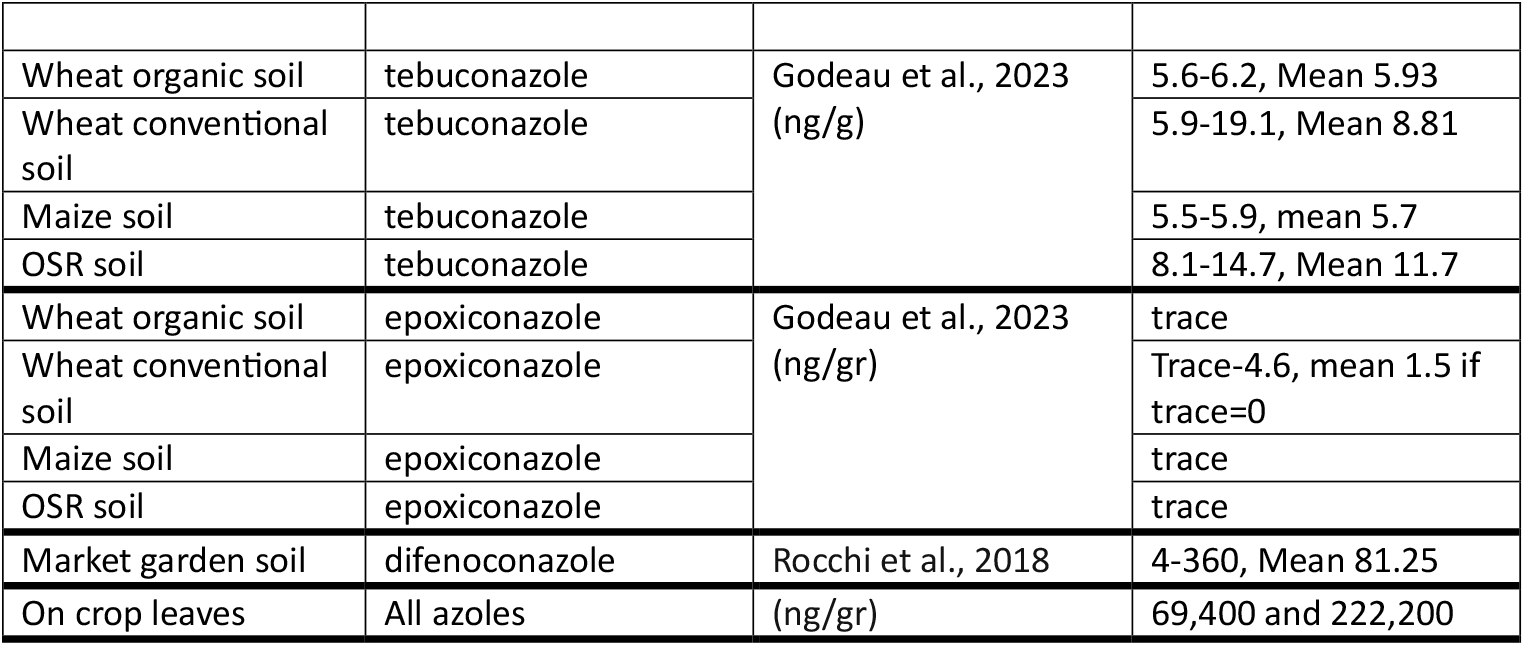

## Supporting information S5

Values for the curvature parameter of the dose-response

Supporting information S5 Values for the curvature parameter of the dose-response
| Fungicide | Pathogen | Curvature | Source |
| --- | --- | --- | --- |
| tricyclazole | *Magnaporthe oryzae* | 2.4 | Kunova et al., 2014 |
| azoxystrobin | *Magnaporthe oryzae* | 0.4 | Kunova et al., 2014 |
| mefenoxam | *Globisporangium species* | 1.6 | Pradhan 2017 |
| cyazofamid | *Pythium aphanidermatum* | 1.67 | Flores & Garzon 2013 |
| propamocarb | *Pythium aphanidermatum* | 1.37 | Flores & Garzon 2013 |
| propiconazole | *Rhizoctonia zeae* | 0.54 | Flores & Garzon 2013 |
| imazalil | *Colletotrichum coccoides* | 1.31 | Uribe & Loria 1994 |
| carbendazim | *Fusarium graminearum* | 0.49 | Gxashekaa et al., 2021 |
| metalaxyl | *Fusarium graminearum* | 0.70 | Gxashekaa et al., 2021 |
| azoxystrobin | *Fusarium graminearum* | 1.11 | Gxashekaa et al., 2021 |
| carbendazim | *Fusarium oxysporum* | 1.68 | Gxashekaa et al., 2021 |
| difenoconazole | *Fusarium oxysporum* | 0.62 | Gxashekaa et al., 2021 |
| tebuconazole | *Pseudocercospora fijiensis* | 1.53 | García-Munguía1 2022 |
| fenbuconazole | *Colletotrichum acutatum* | 2.27 | Mondal et al., 2005 |
| fenbuconazole | *Elsinoe fawcettii* | 1.54 | Mondal et al., 2005 |
| fenbuconazole | *Mycosphaerella citri*, | 0.93 | Mondal et al., 2005 |
| azoxystrobin | *Colletotrichum acutatum* | 1.51 | Mondal et al., 2005 |
| azoxystrobin | *Alternaria alternata* | 1.54 | Mondal et al., 2005 |
| azoxystrobin | *Elsinoe fawcettii* | 1.67 | Mondal et al., 2005 |
| azoxystrobin | *Mycosphaerella citri*, | 0.79 | Mondal et al., 2005 |
| pyraclostrobin | *Colletotrichum acutatum* | 1.51 | Mondal et al., 2005 |
| pyraclostrobin | *Alternaria alternata* | 0.78 | Mondal et al., 2005 |
| pyraclostrobin | *Elsinoe fawcettii* | 1.49 | Mondal et al., 2005 |
| pyraclostrobin | *Mycosphaerella citri*, | 1.02 | Mondal et al., 2005 |

## Supporting information S6

fungicide sensitivity metrics

This section describes sensitivity metrics and how they might be interpreted in relation to resistance risk. Terminology is not always used consistently in the literature, so other interpretations are possible.

### Minimum Inhibitory Concentration (MIC)

MIC is the lowest concentration of an anti-microbial agent which prevents visible growth of the test strain or organism (EUCAST, 1998; Kowalska-Krochmal & Dudek-Wicher, 2021; Mueller et al 2004) under strictly controlled *in vitro* conditions for a standard experimental duration and a standard inoculation density.

The figure illustrates the definition using the log-logistic dose response curve.

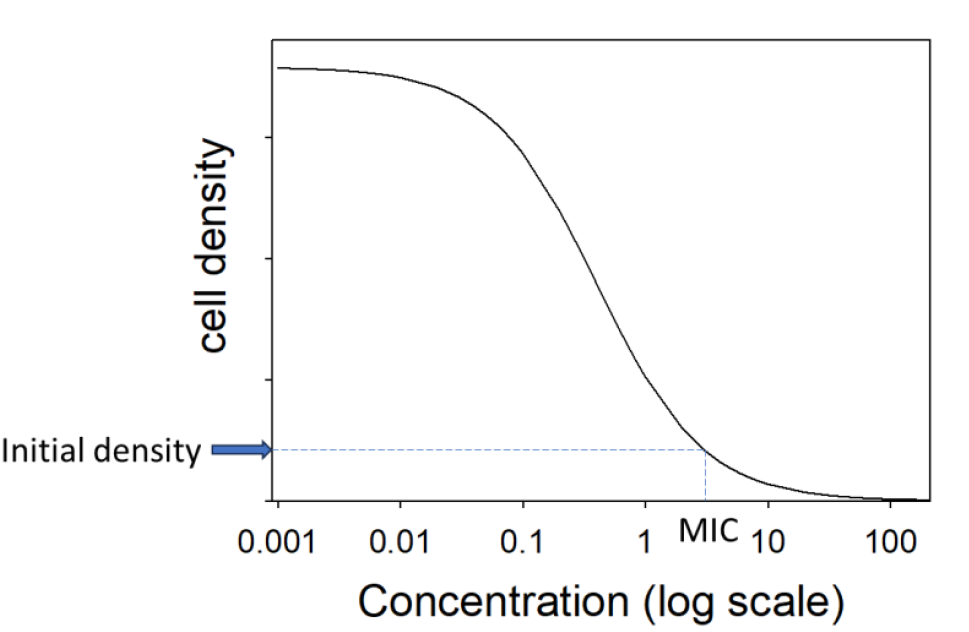

The MIC is usually determined *in vitro* using a nutrient solution or gel with the antifungal added. In soils and compost the energy/nutrient source for fungi is organic matter and the half-life of the fungicide may also be affected by the substrate. Hence the *in vitro* MIC may not represent the minimal inhibitory concentration in practical situations.

In medicine, models are used to assess the value of MIC to determine medication strategies. They include the dynamics of the antimicrobial and the dynamics of the pathogen. Using these models the time that the antimicrobial concentration is above the MIC is determined. Then it is determined what this means for the microbe. Next multiple applications of the antimicrobial are added and a patient treatment scheme is developed. Mueller et al. (2004), Nielsen et al. (2017), Khan et al. (2015), Salas (2020). No models exist to do a similar analysis of the dynamics of the antimicrobial in the soil/compost with a dynamic model for the microbial population. Our model currently assumes a constant antimicrobial concentration, that MIC in the model is the fungicide concentration where *R*_0_=1, which is the same as *r*=0 and that measured concentrations are bio-active (in reality, binding to soil may reduce the bio-active fraction; Reido et al., 2023).

MIC is different from the concentration below which no emergence and selection takes place (NEC, see below). Above the MIC the density of the pathogen (in the lab experiment) does not change. This may seem to suggest that no emergence and selection will take place at and above MIC.

However, at the MIC the total population may be in balance between reproduction/multiplication and death. Above the MIC the total population decreases, but there may still be reproduction, so there could still be selection and emergence above the MIC.

### NOEC or NEC and PNEC

NOEC or NEC: No Effect Concentration. This is usually measured for acute or chronic exposure in the lab or other experimental setting. Chronic exposure is more relevant to emergence and selection that occur over long time periods. The NEC is the concentration below which no detectable effects occur and it often used to denote the absence of adverse effects on non-target organisms. The same limitations of translating a lab NEC to the field apply as described for the MIC (see above).

The PNEC is the concentration of a substance below which an unacceptable effect (on non-targets) is unlikely to occur in practice. The way PNEC is calculated varies, but in most cases a set of NECs resulting from different experiments is used. Either the smallest NEC, or the right-hand limit of the left-hand confidence interval is used. Then the PNEC is calculated by dividing the selected NEC by a factor which depends on how closely the model system relates to the case of interest.

In principle, some effect on emergence and selection can occur at concentrations down to the NEC or PNEC, although the effects may be very small.

### The effect of fitness cost

If there is a fitness cost associated with resistance (for example due to an adverse effect of a resistance mutation on enzyme function) then dose response curves for the resistant and sensitive strains may look like those in the figure below. Below what we term the Minimum Selection Concentration, the sensitive strain will outcompete the resistant strain. This leads to the areas of no selection or emergence in Figs. 7 and 8 in the main text.

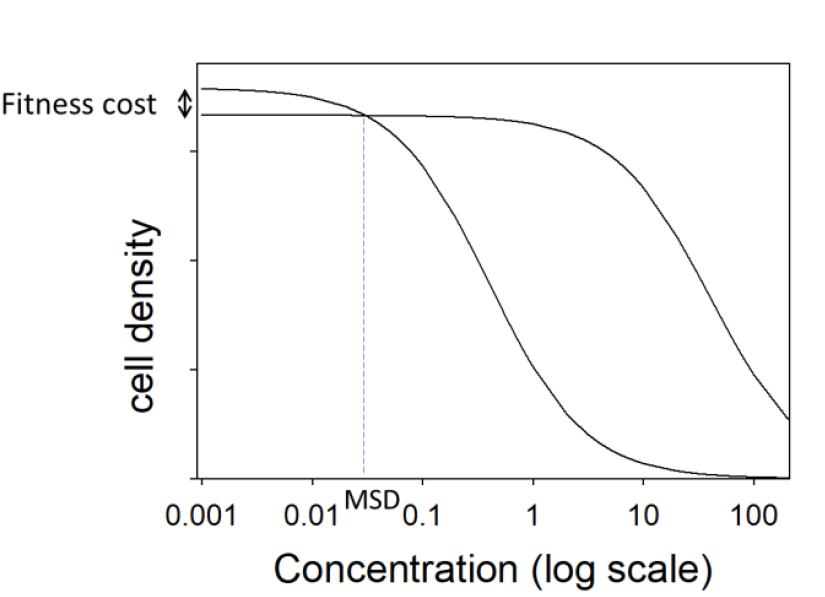

